# geneML: Gene annotation across diverse fungal species using deep learning

**DOI:** 10.64898/2026.05.18.725946

**Authors:** Lisa Vader, Colin J.B. Harvey, Tilmann Weber, Lawrence S. Hon

## Abstract

Accurate gene prediction remains a major bottleneck in fungal genomics, where lineage diversity and alternative splicing challenge existing *ab initio* methods. Here, we present geneML, a deep learning-based gene prediction tool tailored to fungal genomes. Across nine reference genomes spanning diverse fungal taxa, geneML improved gene-level F1 score from 64.9 to 67.1 compared to BRAKER3 with protein-based hints, driven by substantially higher recall (69.0 vs. 64.1) at equivalent precision. geneML also remains fast, averaging around 6 minutes per genome on a standard 8-core CPU. A key feature of geneML is its ability to predict alternative transcripts. Compared to *Fusarium graminearum* Iso-Seq control data, it achieves 41.1% transcript recall and 71.1% precision, outperforming AUGUSTUS (33.8% recall, 48.9% precision), one of the few tools that support isoform prediction. The predicted transcript diversity is consistent with experimentally observed fungal alternative splicing patterns. Reannotation of the curated training dataset further suggests improved biological completeness, with geneML recovering 15.3% more genes containing complete PFAM domains than the reference annotation. These results demonstrate that geneML enables faster, more sensitive, and more biologically informative fungal genome annotation. geneML is available as an open-source command-line tool at https://github.com/hexagonbio/geneML.

**Key Points:**

- geneML improves gene prediction accuracy over both classical and recent deep learning-based methods, while substantially improving recall.
- geneML predicts alternative transcripts with higher precision and recall than AUGUSTUS, expanding functional annotation.
- Runtime was 32-fold decreased over BRAKER3, enabling efficient high-throughput genome annotation.
- geneML identifies novel genes and recovers missing annotations, especially in under-annotated non-Ascomycete genomes.

## Introduction

The annotation of gene locations is foundational to the analysis of fungal genomes. However, current fungal gene prediction methods are lacking in accuracy, completeness and speed. Eukaryotic genomes are difficult to annotate accurately due to their complex gene structures, which often include multiple introns, overlapping genes, and multiple alternative transcripts per locus.^1^ Experimental evidence such as RNA sequencing (RNA-seq) has improved the quality of eukaryotic gene predictions.^2^ However, acquiring RNA-seq data is both costly and experimentally demanding. It also has technical limitations; lowly expressed or conditionally expressed genes may go undetected, and reconstructing transcripts from short-read data is prone to errors.^3^ Therefore, there is a need for *ab initio* gene prediction that provides reliable, fast, and reproducible eukaryotic genome annotation without relying on RNA-seq data.

One of the oldest and most widely used tools for *ab initio* eukaryotic gene prediction is AUGUSTUS. AUGUSTUS predicts genes using a generalised Hidden Markov Model (HMM) trained on species-specific parameters, which captures the gene structure, splice sites and codon usage typical of the species of interest.^4^ AUGUSTUS was shown to outperform other HMM based gene predictors in a recent benchmarking study across 147 eukaryotic species.^5^ The performance of AUGUSTUS depends on high-quality species-specific gene models. Training such models requires manual curation of a reference annotation, which is time-consuming, difficult to standardise, and limited by the availability of reference data. The BRAKER pipeline overcomes this by automating gene model training, removing the need for manual intervention. It runs unsupervised gene prediction with GeneMark^6^ and uses these preliminary predictions for training an AUGUSTUS model, which is used to generate the final gene predictions. It is also possible to use RNA-seq data and/or homology evidence from protein databases as additional evidence for gene prediction.^2^

Recently, there has been a growing interest in gene prediction methods that are independent of species-specific gene models.^1^ Because these models rely on pre-existing annotations, they intrinsically perform better for well-characterised species than for poorly-studied or novel organisms. Additionally, since they rely on species-specific gene structure, they fail to detect atypical genes, such as genes acquired by horizontal gene transfer, and are inherently unsuitable for metagenomic annotation.

Various tools have been developed that apply deep learning to eukaryotic gene prediction. These deep learning models are designed to recognise many different gene structures regardless of the species background, thereby avoiding the core limitations of approaches based on species-specific models. Additionally, deep learning models can offer substantial computational advantages over traditional approaches. Convolutional neural networks (CNNs) are especially suited to the task of gene prediction, as they are able to detect motifs (e.g. splice sites, start and stop codons, transcription factor binding sites) regardless of where they appear in the sequence window, and identify long-range dependencies between motifs.^7,8^

Current deep learning approaches for eukaryotic gene prediction include Helixer^9^, which employs a hybrid CNN and Long short-term memory (LSTM) architecture followed by an HMM-based postprocessing step, and ANNEVO^10^, which uses a mixture-of-experts model based on CNNs followed by a Viterbi decoding algorithm. Both of these tools show promising results compared to traditional approaches. Compared to AUGUSTUS, Helixer shows higher sensitivity but lower precision^1^, while ANNEVO reports higher sensitivity and precision compared to both AUGUSTUS and BRAKER3 in its original publication.

While Helixer and ANNEVO support fungal gene prediction, neither is designed specifically for fungi, potentially limiting coverage of this lineage. Furthermore, existing deep learning gene prediction tools output a single transcript per locus and cannot predict alternative transcripts. Alternative transcripts are multiple distinct mRNA isoforms that are produced from the same locus. Previously, alternative transcripts were thought to be rare in fungi, but they are now recognised as widespread mechanisms for regulation of growth, development and environmental adaptation, with important roles in virulence and drug resistance.^11–14^ Accurately annotating these alternative transcripts is therefore biologically important, particularly for studying fungal pathogenesis.^12,15^

Here we present geneML, a deep learning-based gene predictor trained on a large dataset of fungal genomes, which outperforms existing tools in whole-gene prediction accuracy and runtime, while also predicting alternative transcripts that we show correspond to real transcripts.

## Methods

### Training set generation

We collected all fungal genome records containing gene annotations available at NCBI at December 19th 2025 13:41 UTC (n=6381). We excluded genomes with a contig N50 below 50 kbp, genomes with associated warnings, and genomes with a gene density lower or higher than expected (below 200 or over 1000 genes per Mbp). We selected one genome per genus, prioritising genomes present in RefSeq and subsequently selecting the genome with the highest contig N50. This resulted in a dataset containing 761 genomes; 479 within Ascomycota, 189 within Basidiomycota, 43 within Mucoromycota, 27 within Chytridiomycota, 11 within Microsporidia, 7 within Zoopagomycota and 5 within Blastocladiomycota (Table S1).

From these genomes, we extracted annotated coding sequence (CDS) regions with a valid gene structure (start codon, stop codon, in frame, and no internal stop codons) including flanking context. We excluded regions annotated as pseudogenes and mitochondrial genes. In order to prevent large genomes from dominating the training set, we only included up to 10,000 regions per genome. For multi-transcript loci, we included annotated CDS sequences as separate entries.

We annotated each position in the sequence with a single class label, one of: *CDS start, CDS end, exon start, exon end, is exon, is intron, intergenic*. In order to acquire fixed length training inputs, we split each sequence into chunks of twice the context length of the model, using padding with Ns where necessary.

### Model architecture and training

We designed a CNN architecture to predict the presence of a gene and its structure given a genomic sequence. We leveraged the SpliceAI^8^ residual deep learning architecture, which can predict splice sites in human pre-mRNAs over long sequence contexts, and adapted it for fungal gene calling. Since fungal genes are typically much smaller than human genes, we reduced the number of convolutional layers to 10, thereby significantly reducing the model size. The geneML CNN architecture consists of four regular convolutional layers, four dilated layers with atrous rate 4, and two dilated layers with atrous rate 10, all with a fixed kernel size of 11, yielding a total receptive field of 800 bp. We used a 7-class softmax classification head that predicts the gene annotation class for each base in a sequence.

The model was trained using the Adam optimizer^16^ with learning rate warmup, on a fixed 90/10 train/test split. Model performance was assessed on the held-out test set using top-1 accuracy (the fraction of true sites recovered when predicting an equal number of top-scoring candidates) averaged across the *exon start* and *exon end* classes, which were the classes most impactful for downstream gene calling accuracy. Training continued until top-1 accuracy reached a plateau, with a patience of two epochs, upon which the learning rate decayed and training resumed for a maximum of two such decay steps.

### geneML

The geneML gene prediction pipeline consists of two steps: an inference step and a gene calling step (Figure 1).

**Figure 1:**
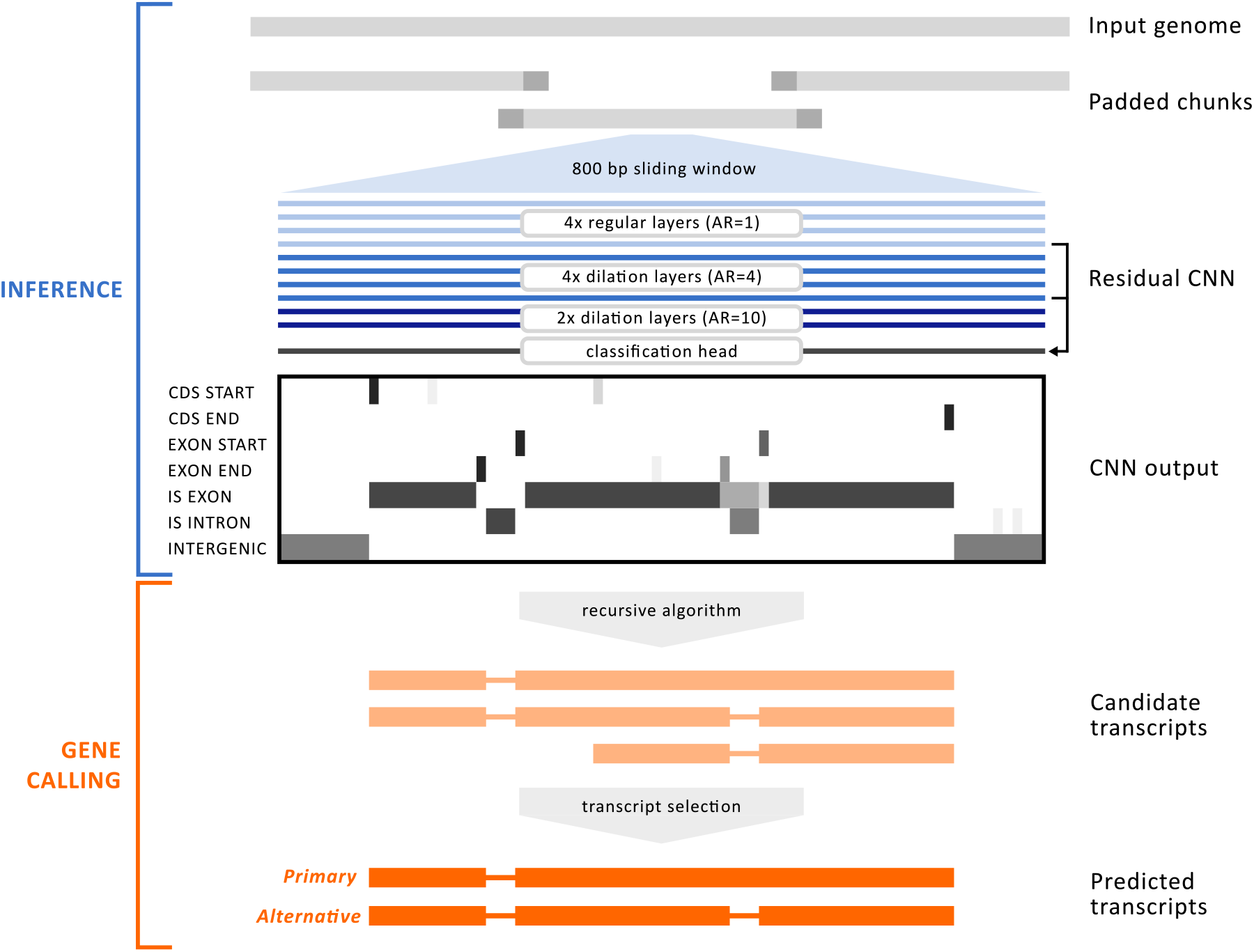
Overview of the geneML pipeline for fungal gene prediction, consisting of an inference step and a gene calling step. During CNN inference, the input genome sequence is split into padded chunks, and a sliding window approach is used to pass 800 bp segments to a dilated residual CNN for per-nucleotide classification into 7 classes. During gene calling, candidate transcripts are built from the CNN output using a constrained depth-first recursive algorithm. The most likely transcript is selected based on length, exon/intron score agreement and the confidence of predicted splice sites and coding sequence boundaries. Where there are additional variants with high-confidence boundary scores, multiple transcripts are predicted. The transcript with the longest CDS is labeled as the primary transcript. CNN = Convolutional Neural Network, CDS = Coding Sequence.

During inference, geneML’s pretrained CNN receives a DNA sequence and predicts per-position probabilities for each of the six annotation classes it was trained on (*CDS start, CDS end, exon start, exon end, is exon* and *is intron*). In order to control memory usage, very large sequences are split into overlapping chunks, where the overlap provides the sequence context necessary for accurate predictions at chunk boundaries.

Next, geneML employs a gene calling algorithm to derive the most likely gene structures from the probabilities outputted by the model. We use a depth-first recursive algorithm, which lends itself well to finding multiple valid solutions and thus suits our aim of detecting alternative transcripts.

First, we identify candidate gene-containing regions by looking for a CDS start signal followed by consistently high *is exon* and/or *is intron* probabilities (> 0.2), in the absence of high *is intergenic* probabilities (> 0.8). For each of these gene regions, we apply depth-first recursion to find possible gene calls, based on predicted gene events (CDS start, CDS end, exon start and exon end) within the region. Gene events include all positions where the probability score for one of the relevant classes exceeds both a global minimum threshold and a percentile-based cutoff for the region. During recursion, we prune any branches that violate a set of biological validity constraints, which ensures the quality of gene predictions and dramatically reduces time spent on recursion. These constraints include: starting with a start codon, ending with a stop codon, containing no premature stop codons, having a length divisible by three, having a higher *is exon* signal than *is intron* signal within exons, and having intron and exon sizes within predefined boundaries.

There typically exist many possible overlapping gene calls for a given region with varying degrees of model support. We hypothesize that the gene calls with the strongest model support are most likely to represent true gene structures. Multiple gene calls with distinct structure and strong model support indicate the presence of alternative transcripts. We select the most likely transcript(s) per locus based on a variety of metrics, of which the gene call quality score is the most important one. The quality score is based on two equally weighted metrics: the event consistency score, which captures how well the suggested exons and introns match the model’s *is exon* and *is intron* probabilities, and the border score, which is the average probability for each gene event included in the gene call. The quality score ranges from 0 to 1, with a higher score indicating higher confidence.

For each region, gene calls scoring more than 0.2 below the maximum score are filtered out. We then build an initial selection containing a seed gene call, which is the highest-scoring gene call at the lowest start position, and any gene calls that are valid alternatives to it. A gene call is considered a valid alternative to another if the scores of added events are substantially higher than those of any removed events, or if there is a single event difference with a high absolute model score. From this initial selection, we define the gene call with the longest CDS as the primary transcript. We define the other gene calls as alternative transcripts, and classify the transcript variant based on comparison of gene structure to the primary transcript.

After transcript selection, we remove low-confidence transcripts based on their quality score. For a given genome, quality scores typically follow a bimodal distribution, with the first peak representing short, low-confidence transcripts and the second peak representing longer, high-confidence transcripts (Figure S1). We define a minimum score cutoff based on the local minimum between the two peaks, which filters out the low-confidence transcripts. The optimal cutoff varies depending on the type of genomic data supplied, and this dynamic cutoff approach typically achieves better results than a static cutoff. Users may also supply a custom quality score cutoff in place of the default.

### Benchmarking

We selected a benchmarking dataset of nine high quality genomes which cover much of the diversity within the known fungal taxonomy, representing nine classes within the fungal phyla Ascomycota, Basidiomycota and Glomeromycota (Table S2). All selected reference genome assemblies are considered complete (assembly is at either chromosome or complete level). In order to minimise bias towards a certain annotation method, we selected genomes that have been annotated using a variety of gene prediction methods. All methods utilised transcript evidence in the form of RNA-seq data, expressed sequence tags (ESTs) and/or full length complementary DNA (cDNA) sequences. The majority of methods also applied protein evidence from publicly available protein datasets or experimentally generated proteomics datasets (Table 1).

**Table 1:**
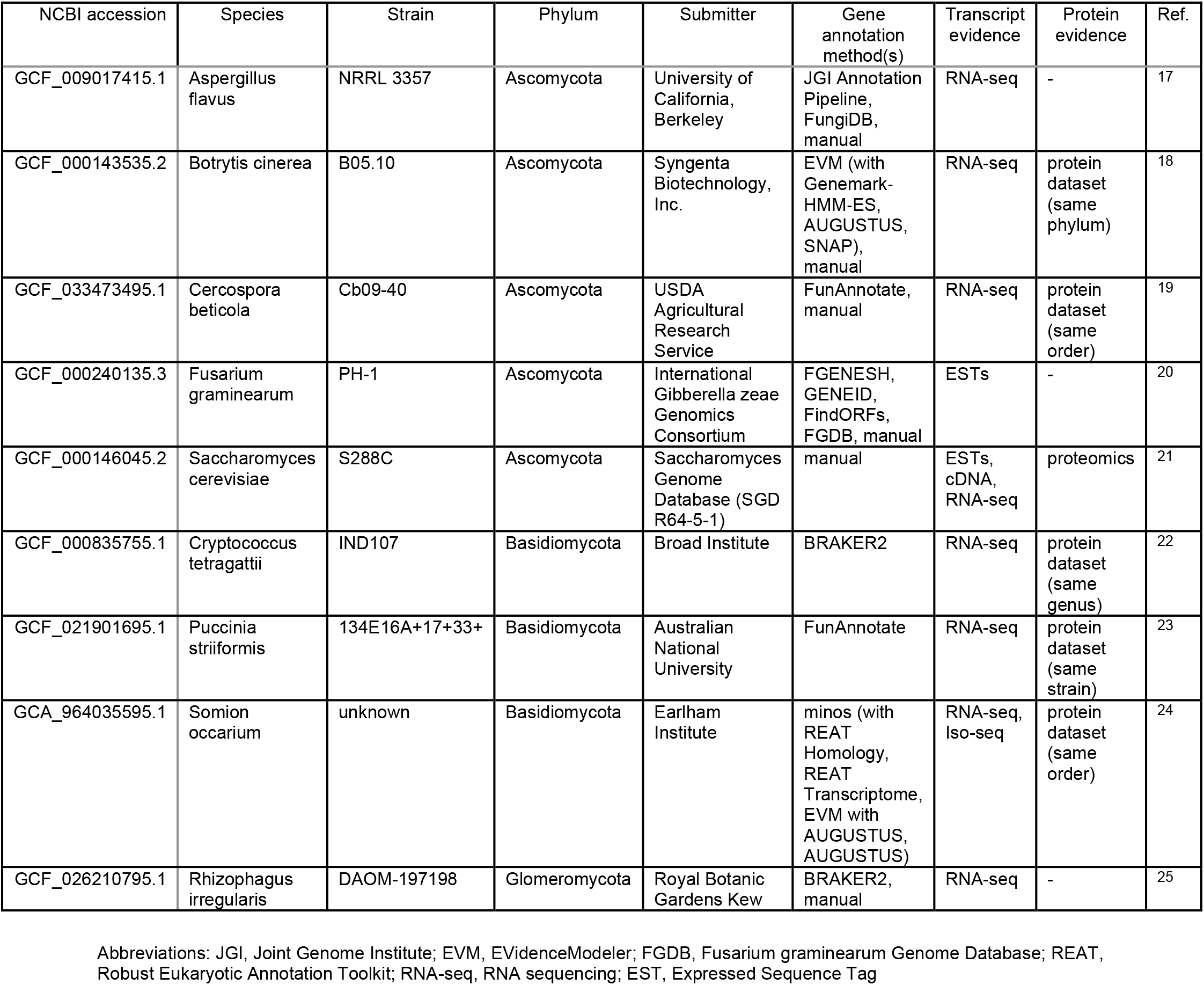
Reference genomes used for benchmarking, including taxonomy, assembly accession, and annotation methodology.

Since we are only interested in evaluating the prediction of nuclear genes, mitochondrial contigs were removed from the genome assembly and annotation files. The gene prediction tools we evaluated do not include untranslated regions (UTRs) in their predictions (with the exception of Helixer). Therefore, UTRs were removed from the reference and Helixer annotations, so that gene and mRNA features only cover the CDS regions.

For each genome, we generated *de novo* gene predictions using geneML v1.1.0, AUGUSTUS v3.5.0^4^, BRAKER3 v3.0.8^2^, Helixer v0.3.6^9^ and ANNEVO v2.2.2^10^. We ran geneML with a model trained on a dataset excluding all benchmarking genera (n=752). We ran AUGUSTUS with the gene model of the species most closely related to the input species (Table S3). We ran BRAKER3 with a protein database that includes fungal sequences in OrthoDB v12.2^26^, any sequences from benchmarking genera removed. We evaluated the performance of each tool by comparing their gene predictions to the reference annotations using gffcompare^27^.

We calculated Recall and Precision at the nucleotide, exon and gene levels. A nucleotide level prediction is considered a True Positive (TP) if the predicted exonic nucleotide overlaps with any exonic nucleotide in the reference. An exon prediction is considered a TP if its start and end coordinates match exactly with a reference exon. A TP gene prediction contains at least one transcript which exactly matches a reference transcript (same start, end and splice site coordinates).

### Validation of alternative transcripts

We validated geneML alternative transcript predictions by comparing to a *Fusarium graminearum* alternative transcript dataset generated by IsoSeq sequencing.^28^ We compared geneML and AUGUSTUS annotations to the Isoseq CDS annotations (UTRs removed) at the transcript level (as previously described). The *F. graminearum* YL1 genome and IsoSeq transcripts were downloaded from http://fgbase.wheatscab.com/Download.

### PFAM domain analysis

We extracted protein sequences using gffread^27^ and identified conserved protein domains based on Pfam 38.2^29^ using HMMER^30^ hmmscan. Domain hits were classified as complete or partial based on pHMM coverage of at least 80%. To avoid false positive partial domain hits, we excluded proteins with a length of less than 100 aa. We mapped Pfam domains to GO terms using Pfam2GO and assigned them to higher-level functional bins using a custom mapping (Table S4). Because mobile genetic element-associated functions are underrepresented in GO, proteins containing characteristic Pfam domains (e.g., integrases, phage structural proteins) were additionally assigned to a ‘Mobile genetic elements’ bin.”

Tool versions, run parameters, and hardware descriptions are provided in Supplementary Methods (Table S5).

## Results

geneML is available as a command-line tool for Linux and macOS and can be installed via pip (pip install geneml). Basic usage requires only a DNA sequence file in FASTA, GenBank or EMBL format. By default, geneML predicts up to five alternative transcripts per locus; this can be restricted to the primary transcript (longest CDS) by setting --max-transcripts 1. The source code and documentation are available at https://github.com/hexagonbio/geneML under a GPL-3.0 license.

### geneML achieves highest gene-level recall and F1 score across diverse fungal genomes

We benchmarked geneML against the HMM based gene prediction tools AUGUSTUS and BRAKER3 and against two deep learning based tools; ANNEVO and Helixer. We compared predicted coding sequence annotations against a dataset of nine high quality reference genomes (Table 1) and calculated recall and precision for each tool (Figure 2, Table 2, Table S6).

**Figure 2:**
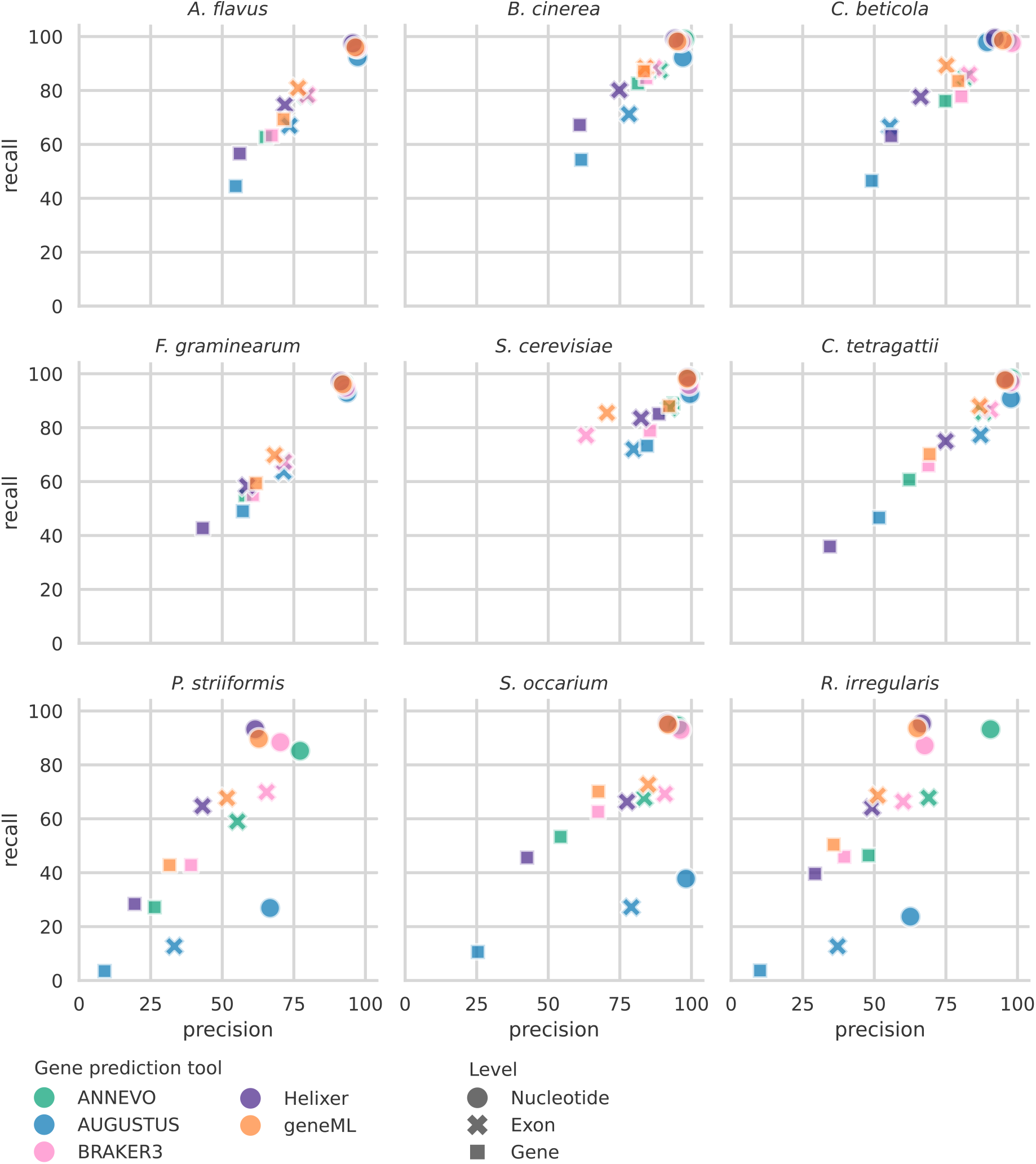
Precision and recall of predicted coding sequence annotations, benchmarked against reference annotations. Precision = Fraction of predicted features that match the reference. Recall = Fraction of reference features recovered. Metrics were computed at the nucleotide, exon, and gene levels using gffcompare.

**Table 2:** Mean recall, precision and F1 score of gene prediction tools across benchmarking genomes. Best performance within a category is shown in bold.

| Level | Tool | Recall (%) | Precision (%) | F1 score (%) |
| --- | --- | --- | --- | --- |
| Nucleotide | ANNEVO | 95.6 ± 4.4 | **93.8 ± 6.8** | **94.7 ± 5.6** |
|  | AUGUSTUS | 71.9 ± 32.1 | 89.1 ± 14.2 | 77.0 ± 26.5 |
|  | BRAKER3 | 94.2 ± 3.9 | 90.6 ± 12.4 | 92.1 ± 8.6 |
|  | Helixer | **97.1 ± 2.1** | 87.5 ± 13.5 | 91.6 ± 8.9 |
|  | geneML | 95.9 ± 2.9 | 88.0 ± 13.9 | 91.4 ± 9.3 |
| Exon | ANNEVO | 76.1 ± 10.8 | **78.8 ± 11.8** | **77.4 ± 11.0** |
|  | AUGUSTUS | 52.2 ± 26.6 | 66.1 ± 19.5 | 56.4 ± 24.4 |
|  | BRAKER3 | 76.6 ± 8.8 | 76.8 ± 12.0 | 76.4 ± 9.4 |
|  | Helixer | 71.6 ± 8.5 | 66.5 ± 13.5 | 68.6 ± 10.7 |
|  | geneML | **79.0 ± 9.2** | 72.1 ± 13.3 | 75.1 ± 10.7 |
| Gene | ANNEVO | 61.4 ± 19.1 | 62.7 ± 19.6 | 62.0 ± 19.4 |
|  | AUGUSTUS | 36.9 ± 24.8 | 44.8 ± 25.1 | 40.0 ± 25.6 |
|  | BRAKER3 | 64.1 ± 14.6 | **65.9 ± 17.3** | 64.9 ± 15.9 |
|  | Helixer | 51.6 ± 17.9 | 47.8 ± 20.5 | 49.5 ± 19.3 |
|  | geneML | **69.0 ± 15.9** | 65.8 ± 20.4 | **67.1 ± 18.4** |

We evaluated gene prediction performance using exact matches at the nucleotide, exon, and gene levels, each representing a progressively more stringent criterion, with performance decreasing accordingly across the different tools. Of these, gene level evaluation is arguably the most biologically informative and meaningful for downstream applications, as it captures protein-level matches.

Consistently across all tools, performance was highest for Ascomycota species and lowest for Basidiomycota and Glomeromycota species, which aligns with the fact that considerably more sequencing data is publicly available for Ascomycota than for other fungal phyla. AUGUSTUS performance was exceptionally low for genomes where no gene model of a closely related species was available (Figure 2, Table S3).

At the nucleotide level, all tools showed high precision (at least 87.5%) and all tools except for AUGUSTUS showed very high recall (at least 94.2%). ANNEVO achieved the highest F1 score (94.7%), which was mostly driven by high precision on *Puccinia striiformis* and *Rhizophagus irregularis*. More substantial differences between tools were found at the exon and gene levels. At the exon level, the highest recall (79.0%) was achieved by geneML, whereas the highest precision (78.8%) and F1 score (77.4%) were reached by ANNEVO. However, at the gene level, geneML achieved the highest recall (69.0%) and F1 score (67.1%), with the highest precision closely tied between BRAKER3 (65.9%) and geneML (65.8%).

For most reference genomes, all tools achieved nucleotide-level precision close to 100%. However, two genomes (*P. striiformis* and *R. irregularis*) show much lower precision than expected. The fact that all gene predictors consistently output more genes than are present in the reference strongly suggests that the reference annotations are incomplete. Therefore, these precision values should be treated with caution. Both genomes show an unusual pattern for ANNEVO, which exhibits higher nucleotide-level precision than the other tools. Interestingly, both the *P. striiformis* and *R. irregularis* reference genomes are present in the training set of ANNEVO. This indicates that the clade-specific models ANNEVO uses are overfitted to these genomes, reproducing reference genes well but with limited ability to predict new genes. Indeed, BRAKER3, Helixer and geneML predict a substantial number of genes with PFAM domain hits which are absent from the reference and ANNEVO annotations (Figure S2, Table S7). geneML identified the highest number of genes containing complete PFAM domains; an increase of 71.9% over the reference annotation for *P. striiformis* and 57.0% for *R. irregularis*.

It is very important to consider that, however unavoidable, the annotations of several reference genomes were (partly) generated by BRAKER2 or other pipelines containing AUGUSTUS (Table 1), which inherently favors these tools. And, as already observed, the deep learning models of ANNEVO and Helixer have been trained on some of the benchmarking genomes, which risks overfitting to the benchmarking set. Sequences from any genus present in the benchmarking dataset were explicitly removed from the geneML benchmarking model. Despite these disadvantages, geneML still outperforms all other tools at gene level recall and F1 score. The fact that these results are consistent in a highly diverse phylogenetic background sets geneML apart as a robust gene prediction tool with exceptional sensitivity for fungal genes.

### geneML is faster than other tools

We compared the runtime of the benchmarked tools across all reference genomes (Table 3, Table S8). When run on 8 CPU cores, geneML was the fastest tool, taking at minimum 1.4 minutes (*Saccharomyces cerevisiae)* and at maximum 12.3 minutes (*R. irregularis*) per genome, with a mean runtime of 6.3 minutes. The second fastest tool was Helixer with a mean runtime of 11.5 minutes per genome. BRAKER3 was the slowest gene prediction tool with a mean runtime of 3.3 hours and took 9.0 hours for *R. irregularis*. geneML outperforms other tools in terms of speed for three reasons: 1) the small size of the CNN, which allows for very fast model inference, 2) an efficient recursive gene calling algorithm relying on depth-first traversal with pruning, and 3) just-in-time compilation using numba^31^, which accelerates performance-critical code paths, but otherwise maintains the readability of Python.

**Table 3:** Mean runtime of each gene prediction tool across benchmarking genomes.

| Tool | Runtime (minutes) | Runtime (hours) |
| --- | --- | --- |
| ANNEVO | 16.40 ± 13.46 | 0.27 ± 0.22 |
| AUGUSTUS | 63.12 ± 65.63 | 1.05 ± 1.09 |
| BRAKER3 | 200.63 ± 137.41 | 3.34 ± 2.29 |
| Helixer | 11.55 ± 9.40 | 0.19 ± 0.16 |
| geneML | **6.25 ± 4.39** | **0.10 ± 0.07** |

All tools exhibited significant positive scaling between runtime and genome size (Figure S3, Table S9), with sub-linear scaling indicating relatively improved efficiency for larger genomes and super-linear scaling indicating the opposite. AUGUSTUS showed super-linear scaling (b = 1.25, r = 0.99), indicating reduced efficiency as genome size increases. ANNEVO (b = 0.97, r = 1.00) and Helixer (b = 0.93, r = 0.99) scaled approximately linearly. BRAKER3 (b = 0.74, r = 0.96) and geneML (b = 0.72, r = 0.74) exhibited sub-linear scaling, suggesting comparatively better performance on larger genomes. Interestingly, geneML showed the weakest correlation between runtime and genome size. This can be explained by the fact that the most time consuming step in geneML is gene calling, which is more strongly influenced by gene complexity than by absolute genome size. For example, annotation of the *Somion occarium* genome, which contains highly complex genes of sometimes more than 40 exons, took longer than expected based on genome size.

### geneML reliably annotates alternative transcripts

A unique feature of geneML is its ability to annotate and classify alternative transcripts. Transcript variation can originate from four sources: alternative transcription start sites (ATSSs), alternative translation initiation (ATI), alternative splicing (AS) and alternative polyadenylation. Of these, the first three have the potential to alter the coding sequence (CDS) of the transcript. GeneML identifies alternative transcripts with CDS-level differences and classifies them into six main types: alternative first exon, alternative last exon, intron retention, exon skipping, alternative 3’ splice site, and alternative 5’ splice site. Transcripts involving more than one of these variations are classified as complex variants.

We compared geneML alternative transcript predictions with a *Fusarium graminearum* reference transcript dataset based on IsoSeq sequencing.^28^ In total, geneML was able to recall 41.2% of transcripts in the IsoSeq dataset, with a precision of 71.1% (Table 4). We also generated transcript predictions with AUGUSTUS, which is the only other tool in our benchmark that explicitly supports alternative transcript predictions. AUGUSTUS recovered fewer transcripts than geneML (33.8%) and at much lower precision (48.0%). For each of the predicted alternative transcript types, we found examples of geneML predictions matching the IsoSeq transcripts (Figure 3, Figure S4).

**Table 4:** Precision and recall of geneML and AUGUSTUS based on *Fusarium graminearum* PH-1 reference transcripts.

| Tool | Recall (%) | Precision (%) |
| --- | --- | --- |
| geneML | **41.2** | **71.1** |
| AUGUSTUS | 33.8 | 48.0 |

**Figure 3:**
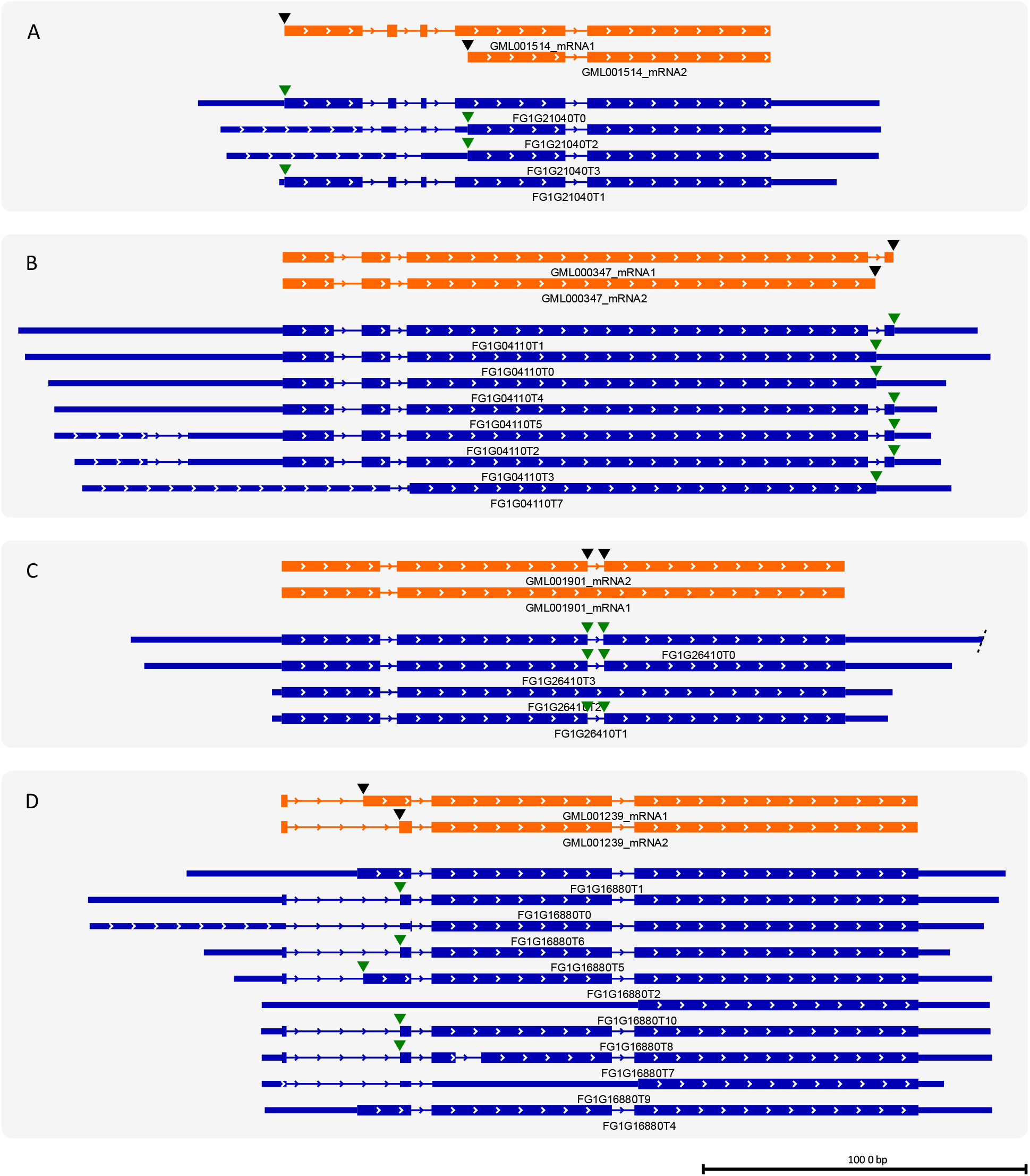
Examples of alternative transcript predictions supported by IsoSeq sequencing data. The four most common transcript variants are shown: A) alternative first exon, B) alternative last exon, C) intron retention, D) alternative 3’ splice site. Transcripts predicted by geneML are shown in orange (predictions do not include UTRs), reference transcripts are shown in dark blue. Gene structures follow IGV conventions: thin lines = introns, medium bars = UTRs, thick bars = coding exons. Black inverted triangles indicate locations of predicted transcript variation, green inverted triangles indicate matching reference variants. *Fusarium graminearum* PH-1 transcript annotations were downloaded from http://fgbase.wheatscab.com/Download.

We also analysed the frequency of alternative transcript predictions among benchmarking genomes (Figure 4A). The most commonly predicted transcript variant was alternative first exon, which was observed at 11-18% of loci across genomes, with a mean value of 13.9%. This truncated transcript variant can be caused by ATSS usage, ATI or by exon skipping or intron retention during AS. In yeast, Translation Initiation Site (TIS) profiling has shown that 388 loci (7% of all loci) produce truncated protein isoforms due to ATSSs or ATI.^32^ In *Cryptococcus neoformans* and *Cryptococcus deneoformans*, 2401 and 2784 ATSSs with the potential to alter the CDS were found respectively (33% and 35% of total TSSs identified).^33^ Although truncated protein isoforms have also been observed in other fungi, their frequency has not been quantified.^12^

**Figure 4:**
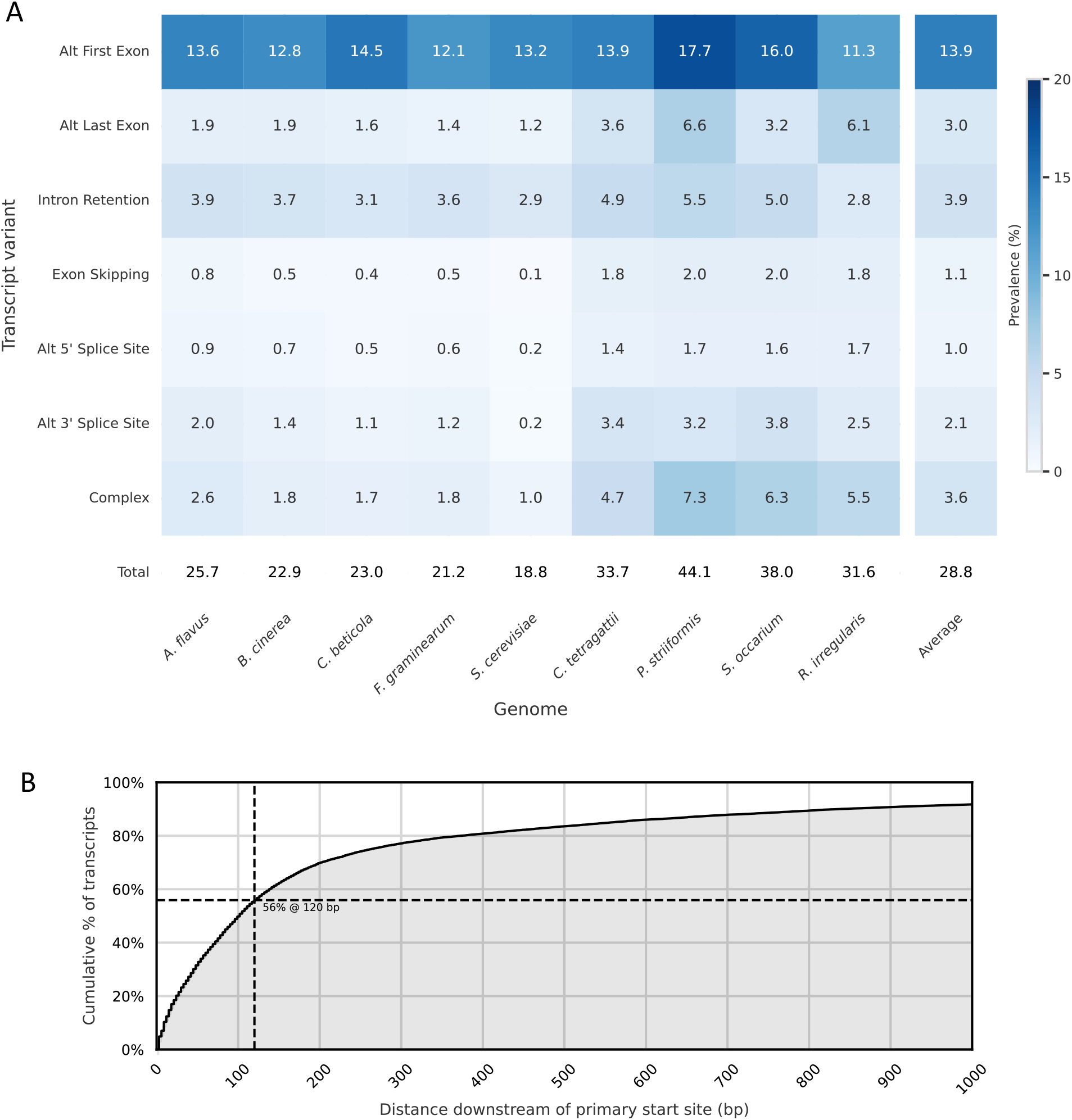
A) Heatmap showing the prevalence of each alternative splicing type per locus as predicted by geneML, for each benchmarking genome. B) Cumulative coverage of alternative start sites by distance from the primary start site. Data includes all predicted alternative transcripts with an altered start site (first exon and complex transcript variants) across benchmarking genomes (n=24,532).

Another observation from the yeast TIS profiling study was that most truncated protein isoforms were close in length to the primary isoform (60% of truncations were within 40 aa from the annotated N terminus). Similarly, for geneML predicted transcripts we observed that 56% of predicted alternative start codons were located up to 120 bp downstream of the primary start codon (Figure 4B). Proximal truncations have been shown to alter the localisation of the produced protein, while the core functional domains remain unchanged.^32,33^

Among transcript variants specific to AS, the most frequently predicted variant was intron retention found at 3. % of loci on average, followed by alternative 3’ splice site (. %), exon skipping (. %) and alternative ‘splice site (. %) (igure A). Intron retention is the most commonly observed type of AS in fungi, followed by alternative 3’ splice site, alternative’ splice site and exon skipping.^11,28^ This matches the frequency of geneML predicted AS events, with the exception of exon skipping, which was reported more frequently than expected. It is challenging to distinguish between real biological variation in exon usage and exons that are simply more difficult for the model to detect, leading to an inflated number of predicted exon skipping events.

The species with most alternative transcript predictions was *P. striiformis*, with 44.1% of loci containing at least one alternative transcript annotation, whereas *S. cerevisiae* had the least alternative transcript annotations at 18.8% of loci (Figure 4A). Most of these (14.4%) were due to alternative first or last exon usage, which corresponds to the fact that ATSS and ATI are common in this species but AS is rare (observed frequency is only 0.2%), as the vast majority of genes do not contain any introns.^11^ However, even though geneML accurately predicts almost all reference genes in *S. cerevisiae* (Figure 2), due to its atypical splice site signatures compared to most fungi, geneML has a tendency to overpredict AS variants in this species.

### geneML reannotation of training genomes produces new valid gene annotations

In the benchmarking dataset, we previously observed that geneML improved incomplete reference annotations in two genomes. To determine whether this observation extends to the training dataset, we reannotated all genomes used for training of the geneML CNN. Annotation with geneML increased the total number of gene annotations from a median of 3241 genes per genome (IQR: 2462-4837) in the original annotation to 3547 genes per genome (IQR: 2683-5032) in the reannotation (Table S10). The median length of predicted proteins was slightly increased from 367 aa (gene length: 1101 nt) to 384 aa (gene length: 1152 nt) (Figure S5). The average BUSCO completeness increased from 91.6% to 93.0%.

We assessed the validity of the predicted genes through the presence of conserved protein domains (PFAM domains). The majority (80.9%) of gene annotations that were added by geneML contained partial or complete hits to PFAM domains indicating that these may indeed be functional (Table S11). Overall, geneML increased the number of genes without PFAM domains with 4.1%, genes with partial PFAM domains with 6.4% and genes with complete PFAM domains with 15.3%. Interestingly, the largest gains in genes with complete domains were observed outside of Ascomycota, with increases of 30.0% in Basidiomycota and 25.4% in other phyla (Figure 5A). These are promising results, indicating that geneML can generate novel and reliable gene predictions in clades which are typically challenging to annotate.

**Figure 5:**
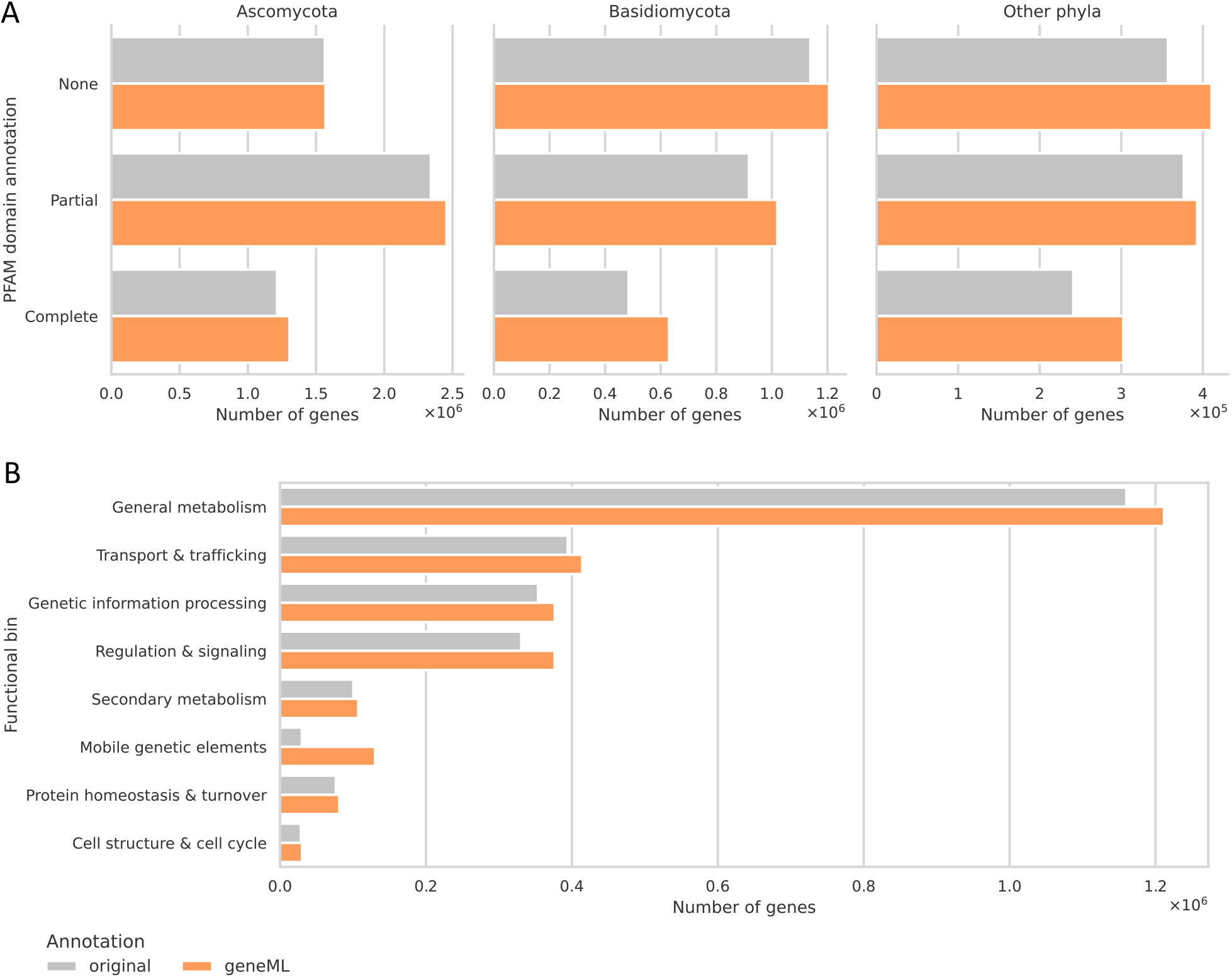
Results of geneML reannotation of training genomes (n=761), compared to original NCBI annotations. A) Total number of annotated genes with complete, partial or no PFAM domain(s), delineated by phylum. A complete domain requires at least 80% profile hidden Markov model (pHMM) coverage. B) Predicted gene functions based on Gene Ontology (GO) annotation of genes. Data only shown for PFAM domains that could be mapped to GO functions and where it was possible to assign a functional bin based on GO.

In order to assess the functional impact of the added gene annotations, we assigned putative functions to genes based on their PFAM annotations. We found that geneML reannotation increased the number of genes in all functional categories (Figure 5B, Table S12). Strikingly, there was a 337.2% increase in genes associated with mobile genetic elements (MGEs), especially genes with domains commonly found in transposable elements (TEs) (Table S13).

Although it is well known that fungal genomes are rich in TEs, they are often underannotated in reference genomes. Standard HMM-based gene prediction approaches lack explicit states for repetitive sequences and tend to produce many spurious gene calls based on fragmented ORFs within repeat-rich regions, which inflate false positive predictions and add significant computational overhead. For this reason, repeats have traditionally been masked in eukaryotic genome analyses. However, it is increasingly appreciated that repetitive DNA carries important biological information and that excluding these regions can limit or bias genomic interpretation.^34^ Because of the nature of TEs, which tend to evolutionarily degrade rapidly, a portion of geneML TE annotations are expected to represent pseudogenes. However, we also identify a significant number of domains associated with domesticated TEs, such as the chromo domain and PEG10/RTL1 protease, as well as intact retrotransposon machinery such as reverse transcriptases and integrase domains, indicative of potentially active LTR retrotransposon lineages (Table S13).

Additionally, annotation with geneML expanded the ‘Regulation & signaling’ bin by 13.8%, particularly through the addition of transcription factor-encoding genes, which are known to be enriched for AS in mammals.^35^ This suggests that our AS-sensitive gene calling method may be especially well suited to identifying transcription factors. We also see significant improvements in the detection of genes with complex structures, such as those encoding protein kinases and Major Facilitator Superfamily transporters (Table S13).

## Discussion

HMM-based methods have long been valuable tools for identifying eukaryotic genes in genomic sequences. Their core strength lies in modeling species-specific gene structures, which yields high specificity. However, this same quality is also a limitation, as it constrains these methods to reproducing known gene structures and reduces their ability to detect novel genes or generalize beyond the genomes of well-characterized species. Meanwhile, recent advances in deep learning architectures have opened new possibilities for gene prediction.

We present a new deep learning-based pipeline for fungal gene prediction, geneML, which outperforms both traditional HMM-based tools and deep learning-based gene predictors in gene-level F1 score (Figure 2), demonstrating especially high recall. Notably, geneML also achieves a substantial speedup over BRAKER3 (Table 3), the next best-performing tool by gene-level F1 score. This combination of high sensitivity and speed makes geneML especially well-suited for high-throughput fungal genome annotation, metagenomic analyses, and rapid annotation of specific genomic regions of interest.

Notably, geneML detects novel genes even in the genomes used for training, demonstrating that rather than memorising training data, it successfully extracts and applies generalizable sequence patterns. The largest annotation improvements were observed in non-Ascomycete phyla, suggesting that reference annotations for these groups are more frequently incomplete, and that geneML is particularly effective at recovering missing genes in these genomes. Functional classification of the added annotations further reveals that geneML improves gene detection across all functional categories, with especially strong performance in transposable element-associated genes, transcription factor genes, and genes with complex structures.

A key novel feature of geneML is the prediction of alternative transcripts. Although alternative transcripts represent an important layer of regulatory complexity in fungal biology, they have largely been overlooked in bioinformatics tools. geneML achieves higher recall and substantially higher precision in alternative transcript prediction than AUGUSTUS, to our knowledge the only other tool supporting this feature. Furthermore, the frequency of predicted transcript variants is consistent with established patterns in fungal biology.

The ability to predict alternative transcripts has significant biological implications. Protein isoforms arising from alternative transcripts frequently differ in domain architecture and subcellular localisation^14^, meaning that analyses restricted to primary isoforms risk underestimating the true functional diversity of a species. Beyond expanding the overall protein repertoire, alternative transcript prediction can help identify genes more likely to be involved in environmental adaptation. For example, alternative splicing is known to be enriched in genes associated with stress responses and pathogenicity, including drug resistance genes.^11,15,36^ In a comparative genomics context, differences in isoform repertoire between related species may further provide valuable insight into divergent adaptive strategies.

The foundation of geneML is a carefully curated training set of fungal genes drawn from more than 750 genera, designed to capture as much fungal diversity as possible. Since the availability of annotated genomes is heavily skewed towards well-studied genera, we reduced sampling bias by restricting the dataset to a single high-quality genome per genus. We also explored training models at phylum resolution, but found it did not improve performance (data not shown). To ensure data quality, we applied strict filtering based on valid protein-coding gene structure, and randomly sampled up to 10,000 valid genes per genome to prevent large genomes from dominating the dataset. Interestingly, despite a large portion of the training set consisting of predicted genes, which inevitably contain errors, geneML successfully extracts generalizable gene structure patterns from this noisy input, ultimately generating predictions that exceed the quality of the training data itself.

The CNN used by geneML is relatively simple in structure compared to other deep learning-based gene callers, which combine CNNs with LSTMs or Transformer layers to capture very long-range sequence context.^9,10^ Instead, geneML employs a residual dilated CNN to capture sequence signatures at short to medium range. Using a receptive field of only 800 bp, the geneML CNN is considerably smaller than those of comparable tools. However, for fungal gene prediction, where introns are typically short, this architecture works well. We considered larger receptive fields, but did not observe performance advantages (data not shown), suggesting that 800 bp is sufficient for capturing fungal introns. As an added advantage, the small CNN size makes it computationally efficient, freeing up resources for a more thorough gene decoding step. Rather than applying Viterbi decoding, which is standard in many gene annotation tools but is limited to returning a single optimal solution, geneML uses a constrained depth-first recursive algorithm that naturally enumerates multiple valid annotation paths, enabling the prediction of alternative transcripts. Efficiency is maintained through two-tier candidate filtering, which limits the number of gene event candidates considered at each step, and through early pruning of paths that violate biological validity constraints. Together, these design choices enable fast and flexible generation of alternative gene predictions.

Despite its strong overall performance, geneML has several limitations. The recursive algorithm is not fully exhaustive; recursions are capped at 100,000 to avoid path explosion, which can in rare cases result in incomplete gene calls. The biological validity constraints also make the algorithm sensitive to sequence errors. For example, a single base pair error can produce an apparent frameshift that disrupts results. As with any machine learning model, performance is expected to decline for fungal clades poorly represented in the training set. Additionally, short single-exon genes are more likely to be missed as they are underrepresented in the training data.

There are several clear directions for future development of geneML. While sampling of genomes across taxonomy implicitly adds gene structure diversity to the training set, explicitly balancing the training set by gene features such as exon number and splice site signatures could improve coverage of less common gene structures. Coverage of rare fungal phyla remains limited by the scarcity of high-quality annotated assemblies. Here, sequence simulation offers a potential route to augmenting training data. Prediction of full-length transcripts including UTRs would be a valuable extension as long-read RNA sequencing becomes more widely accessible. Beyond improvements to the current tool, the CNN’s ability to detect fungal gene structure signatures could be leveraged for classifying contigs in metagenomic datasets. Finally, while geneML was developed for fungal gene prediction, the training set selection procedure and overall gene calling approach generalise well to other datasets, making extension to other eukaryotic lineages or sequence types (such as non-coding RNAs) a natural long-term direction, although this may require adjustments to model architecture and gene calling parameters.

## Acknowledgements

The authors would like to thank Hexagon Bio for providing the resources and environment for advancing this research and for hosting Lisa Vader during her secondment. In particular, we would like to thank Jerry Xu for his help with software development, Brian Naughton, Tara Arvedson, and Maureen Hillenmeyer for supporting this project, and Yuling Liu, Eli Moss, Pablo Cordero and the rest of the Data Team, as well as Joe Spraker, Bruno Perlatti and Anthony DeNicola for helpful feedback. The authors also wish to thank Pablo Cruz-Morales, David Faurdal and Tue Sparholt Jørgensen for their helpful discussions and Thea Laulund Lunden and Mikael Terp for testing geneML.

## Data availability

The data underlying this article are available in Zenodo, at https://doi.org/10.5281/zenodo.20206540.

