## Supplementary Materials for "geneML: Gene annotation across diverse fungal species using deep learning"

### Supplementary Methods

#### Benchmarking

All benchmarks were run on a server with two Intel Xeon Silver 4410Y processors (12 cores each, 24 cores / 48 threads total, up to 3.9 GHz) and 125 GB RAM, running Ubuntu 22.04. Each tool was executed in an isolated Docker container pinned to 8 CPUs and a single NUMA node. Jobs were run sequentially on an otherwise idle machine.

#### BUSCO completeness

We evaluated the completeness of predicted genes using BUSCO based on the longest protein isoform per locus (to avoid false positive duplicates).

### Supplementary Tables

Table S1: Training set details (provided as a separate Excel file)

Table S2: Number of genomes available at NCBI for different fungal phyla and classes. Classes represented in the benchmarking dataset are shown in bold. Data retrieved from the NCBI Datasets Taxonomy Browser (https://www.ncbi.nlm.nih.gov/datasets/taxonomy) at 09/03/2026.

| Kingdom | Phylum | Class | Number of genomes |
|---|---|---|---|
| Fungi | | | 23832 |
| | Ascomycota | | 19452 |
| | | **Dothideomycetes** | 2671 |
| | | **Eurotiomycetes** | 3072 |
| | | **Leotiomycetes** | 431 |
| | | **Saccharomycetes** | 3233 |
| | | **Sordariomycetes** | 5833 |
| | | Other classes | 4212 |
| | Basidiomycota | | 3340 |
| | | **Agaricomycetes** | 1813 |
| | | **Pucciniomycetes** | 151 |
| | | **Tremellomycetes** | 593 |
| | | Other classes | 783 |
| | Mucoromycota | | 532 |

| | | | |
|---|---|---|---|
| | | **Glomeromycetes** | 148 |
| | | Other classes | 384 |
| | Other phyla | | 505 |

Table S3: Models used for running AUGUSTUS.

| Input species | Model name | Shared rank |
|---|---|---|
| Aspergillus flavus | aspergillus_oryzae | species |
| Botrytis cinerea | botrytis_cinerea | species |
| Cercospora beticola | botrytis_cinerea | subphylum |
| Cryptococcus tetragattii | cryptococcus_neoformans_gattii | genus |
| Fusarium graminearum | fusarium_graminearum | species |
| Puccinia striiformis | ustilago_maydis | phylum |
| Rhizophagus irregularis | rhizopus_oryzae | phylum |
| Saccharomyces cerevisiae | saccharomyces_cerevisiae_S288C | species |
| Somion occarium | phanerochaete_chrysosporium | order |

Table S4: Mapping of GO annotations to functional bins

| Bin ID | Bin Name | Biological Process IDs | Molecular Function IDs |
|---|---|---|---|
| 1 | Transposable element–related | GO:0006313 | GO:0003964,GO:0004540, GO:0004523 |
| 2 | Secondary metabolism | GO:0019748,GO:0030639, GO:0019184,GO:0016114 | GO:0004497,GO:0016218, GO:1904091,GO:0010333 |
| 3 | Regulation & signaling | GO:0007165,GO:0050896 | GO:0004672,GO:0016301, GO:0004713,GO:0000155, GO:0000981,GO:0000976, GO:0003700 |
| 4 | Genetic information processing | GO:0006259 | GO:0003676,GO:0003723, GO:0003729,GO:0003735, GO:0003887,GO:0003735 |
| 5 | Protein homeostasis & turnover | GO:0006457,GO:0006508 | GO:0004842 |
| 6 | Transport & trafficking | GO:0006810 | GO:0005215,GO:0022857, GO:0015075,GO:0015291, GO:0008324 |
| 7 | Cell structure & cell cycle | GO:0007010,GO:0051301 | GO:0003779,GO:0008017, GO:0003777,GO:0005198 |
| 8 | General metabolism | GO:0008152 | GO:0003824,GO:0016491, GO:0016787,GO:0016740, GO:0016853,GO:0016791 |

Table S5: Software and databases used in this study. Tool versions, database releases, and non-default parameters are listed.

| Name | Version | Options/Notes |
|---|---|---|
| geneML | v1.1.0 | |
| geneML (benchmarking) | v1.1.0 | --model [benchmarking model], --context-length 800 |
| AUGUSTUS | v3.5.0 | --gff3=on, --species=[x] (Table S3) |
| AUGUSTUS (alternative transcripts) | v3.5.0 | --gff3=on, --alternatives-from-sampling=true, --maxtracks=5, --species=fusarium_graminearum |
| BRAKER3 | v3.0.8 | --fungus, --gff3, --prot_seq=[filtered OrthoDB v12.2] |
| Helixer | v0.3.6 | --lineage fungi |
| ANNEVO | v2.2.2 | --model_path ANNEVO_model/ANNEVO_Fungi.pt |
| gffcompare | v0.12.10 | --no-exon-merge, --strict-match, -e 0 |
| gffread | v0.12.8 | |
| HMMER | v3.3.2 | --cut_tc |
| OrthoDB | v12.2 | Fungal sequences, benchmarking genera removed |
| Pfam | 38.2 | |
| Pfam2GO | v2026/03/02 | https://current.geneontology.org/ontology/external2go/pfam2go |
| BUSCO | v6.0.0 | --mode protein, --lineage fungi_odb12 |

Table S6: Precision, recall and F1 score of predicted coding sequence annotations, benchmarked against reference annotations.

| Tool | Species | Level | Recall | Precision | F1 score |
|---|---|---|---|---|---|
| ANNEVO | Aspergillus flavus | Exon | 78 | 79.2 | 78.6 |
| ANNEVO | Aspergillus flavus | Gene | 62.7 | 65.2 | 63.9 |
| ANNEVO | Aspergillus flavus | Nucleotide | 96.6 | 97 | 96.8 |
| ANNEVO | Botrytis cinerea | Exon | 87.5 | 88.9 | 88.2 |
| ANNEVO | Botrytis cinerea | Gene | 82.6 | 81.3 | 81.9 |
| ANNEVO | Botrytis cinerea | Nucleotide | 99 | 97.7 | 98.3 |
| ANNEVO | Cercospora beticola | Exon | 84.9 | 81.7 | 83.3 |
| ANNEVO | Cercospora beticola | Gene | 76.1 | 74.8 | 75.4 |
| ANNEVO | Cercospora beticola | Nucleotide | 98.7 | 96.7 | 97.7 |
| ANNEVO | Cryptococcus tetragattii | Exon | 85.7 | 88.1 | 86.9 |
| ANNEVO | Cryptococcus tetragattii | Gene | 60.7 | 62.2 | 61.4 |
| ANNEVO | Cryptococcus tetragattii | Nucleotide | 98.3 | 97.9 | 98.1 |
| ANNEVO | Fusarium graminearum | Exon | 67.4 | 71.2 | 69.2 |
| ANNEVO | Fusarium graminearum | Gene | 54.8 | 58 | 56.4 |
| ANNEVO | Fusarium graminearum | Nucleotide | 96.5 | 92.6 | 94.5 |
| ANNEVO | Puccinia striiformis | Exon | 59 | 55.3 | 57.1 |

| | | | | | |
|---|---|---|---|---|---|
| ANNEVO | Puccinia striiformis | Gene | 27.2 | 26.4 | 26.8 |
| ANNEVO | Puccinia striiformis | Nucleotide | 85.2 | 77.2 | 81.0 |
| ANNEVO | Rhizophagus irregularis | Exon | 67.8 | 69 | 68.4 |
| ANNEVO | Rhizophagus irregularis | Gene | 46.4 | 48 | 47.2 |
| ANNEVO | Rhizophagus irregularis | Nucleotide | 93.2 | 90.6 | 91.9 |
| ANNEVO | Saccharomyces cerevisiae | Exon | 87.2 | 92.3 | 89.7 |
| ANNEVO | Saccharomyces cerevisiae | Gene | 89 | 93.7 | 91.3 |
| ANNEVO | Saccharomyces cerevisiae | Nucleotide | 98.5 | 99.5 | 99.0 |
| ANNEVO | Somion occarium | Exon | 67.7 | 83.6 | 74.8 |
| ANNEVO | Somion occarium | Gene | 53.3 | 54.3 | 53.8 |
| ANNEVO | Somion occarium | Nucleotide | 94.6 | 95.2 | 94.9 |
| AUGUSTUS | Aspergillus flavus | Exon | 66.9 | 73.5 | 70.0 |
| AUGUSTUS | Aspergillus flavus | Gene | 44.5 | 54.7 | 49.1 |
| AUGUSTUS | Aspergillus flavus | Nucleotide | 92.4 | 97.3 | 94.8 |
| AUGUSTUS | Botrytis cinerea | Exon | 71.2 | 78.2 | 74.5 |
| AUGUSTUS | Botrytis cinerea | Gene | 54.3 | 61.5 | 57.7 |
| AUGUSTUS | Botrytis cinerea | Nucleotide | 92.2 | 97 | 94.5 |
| AUGUSTUS | Cercospora beticola | Exon | 66.6 | 55.4 | 60.5 |

| | | | | | |
|---|---|---|---|---|---|
| AUGUSTUS | Cercospora beticola | Gene | 46.5 | 49.1 | 47.8 |
| AUGUSTUS | Cercospora beticola | Nucleotide | 97.9 | 89.4 | 93.5 |
| AUGUSTUS | Cryptococcus tetragattii | Exon | 77.2 | 87.1 | 81.9 |
| AUGUSTUS | Cryptococcus tetragattii | Gene | 46.6 | 51.7 | 49.0 |
| AUGUSTUS | Cryptococcus tetragattii | Nucleotide | 90.8 | 97.7 | 94.1 |
| AUGUSTUS | Fusarium graminearum | Exon | 63.7 | 71.4 | 67.3 |
| AUGUSTUS | Fusarium graminearum | Gene | 49 | 57.2 | 52.8 |
| AUGUSTUS | Fusarium graminearum | Nucleotide | 92.9 | 93.6 | 93.2 |
| AUGUSTUS | Puccinia striiformis | Exon | 12.7 | 33.3 | 18.4 |
| AUGUSTUS | Puccinia striiformis | Gene | 3.5 | 8.9 | 5.0 |
| AUGUSTUS | Puccinia striiformis | Nucleotide | 26.9 | 66.7 | 38.3 |
| AUGUSTUS | Rhizophagus irregularis | Exon | 12.7 | 37.2 | 18.9 |
| AUGUSTUS | Rhizophagus irregularis | Gene | 3.7 | 10.1 | 5.4 |
| AUGUSTUS | Rhizophagus irregularis | Nucleotide | 23.7 | 62.6 | 34.4 |
| AUGUSTUS | Saccharomyces cerevisiae | Exon | 72 | 79.8 | 75.7 |
| AUGUSTUS | Saccharomyces cerevisiae | Gene | 73.3 | 84.5 | 78.5 |
| AUGUSTUS | Saccharomyces cerevisiae | Nucleotide | 92.5 | 99.5 | 95.9 |
| AUGUSTUS | Somion occarium | Exon | 27.2 | 79 | 40.5 |

| | | | | | |
|---|---|---|---|---|---|
| AUGUSTUS | Somion occarium | Gene | 10.6 | 25.4 | 15.0 |
| AUGUSTUS | Somion occarium | Nucleotide | 37.8 | 98.1 | 54.6 |
| BRAKER3 | Aspergillus flavus | Exon | 78.3 | 79.8 | 79.0 |
| BRAKER3 | Aspergillus flavus | Gene | 63.3 | 67.3 | 65.2 |
| BRAKER3 | Aspergillus flavus | Nucleotide | 95.8 | 97.2 | 96.5 |
| BRAKER3 | Botrytis cinerea | Exon | 88.3 | 86.9 | 87.6 |
| BRAKER3 | Botrytis cinerea | Gene | 84.6 | 84.2 | 84.4 |
| BRAKER3 | Botrytis cinerea | Nucleotide | 98.1 | 96.5 | 97.3 |
| BRAKER3 | Cercospora beticola | Exon | 85.9 | 83.1 | 84.5 |
| BRAKER3 | Cercospora beticola | Gene | 77.8 | 80.4 | 79.1 |
| BRAKER3 | Cercospora beticola | Nucleotide | 97.6 | 98 | 97.8 |
| BRAKER3 | Cryptococcus tetragattii | Exon | 86.7 | 90.5 | 88.6 |
| BRAKER3 | Cryptococcus tetragattii | Gene | 66 | 68.9 | 67.4 |
| BRAKER3 | Cryptococcus tetragattii | Nucleotide | 96.9 | 97.6 | 97.2 |
| BRAKER3 | Fusarium graminearum | Exon | 67.3 | 71.6 | 69.4 |
| BRAKER3 | Fusarium graminearum | Gene | 55 | 60.7 | 57.7 |
| BRAKER3 | Fusarium graminearum | Nucleotide | 94.8 | 93.1 | 93.9 |
| BRAKER3 | Puccinia striiformis | Exon | 69.9 | 65.5 | 67.6 |

| | | | | | |
|---|---|---|---|---|---|
| BRAKER3 | Puccinia striiformis | Gene | 42.8 | 39.1 | 40.9 |
| BRAKER3 | Puccinia striiformis | Nucleotide | 88.4 | 70.3 | 78.3 |
| BRAKER3 | Rhizophagus irregularis | Exon | 66.4 | 60.1 | 63.1 |
| BRAKER3 | Rhizophagus irregularis | Gene | 45.9 | 39.5 | 42.5 |
| BRAKER3 | Rhizophagus irregularis | Nucleotide | 87.2 | 67.6 | 76.2 |
| BRAKER3 | Saccharomyces cerevisiae | Exon | 77.1 | 63.2 | 69.5 |
| BRAKER3 | Saccharomyces cerevisiae | Gene | 78.9 | 85.5 | 82.1 |
| BRAKER3 | Saccharomyces cerevisiae | Nucleotide | 95.8 | 99.2 | 97.5 |
| BRAKER3 | Somion occarium | Exon | 69.2 | 90.7 | 78.5 |
| BRAKER3 | Somion occarium | Gene | 62.6 | 67.4 | 64.9 |
| BRAKER3 | Somion occarium | Nucleotide | 93 | 96.2 | 94.6 |
| geneML | Aspergillus flavus | Exon | 80.9 | 76.5 | 78.6 |
| geneML | Aspergillus flavus | Gene | 69.3 | 71.4 | 70.3 |
| geneML | Aspergillus flavus | Nucleotide | 96 | 96.5 | 96.2 |
| geneML | Botrytis cinerea | Exon | 88.5 | 83.9 | 86.1 |
| geneML | Botrytis cinerea | Gene | 87.1 | 83.4 | 85.2 |
| geneML | Botrytis cinerea | Nucleotide | 98.2 | 95.1 | 96.6 |
| geneML | Cercospora beticola | Exon | 89.2 | 75.2 | 81.6 |

| | | | | | |
|---|---|---|---|---|---|
| geneML | Cercospora beticola | Gene | 83.5 | 79.3 | 81.3 |
| geneML | Cercospora beticola | Nucleotide | 98.6 | 94.9 | 96.7 |
| geneML | Cryptococcus tetragattii | Exon | 88 | 86.8 | 87.4 |
| geneML | Cryptococcus tetragattii | Gene | 70.2 | 69.3 | 69.7 |
| geneML | Cryptococcus tetragattii | Nucleotide | 97.6 | 95.7 | 96.6 |
| geneML | Fusarium graminearum | Exon | 69.8 | 68.3 | 69.0 |
| geneML | Fusarium graminearum | Gene | 59.4 | 61.8 | 60.6 |
| geneML | Fusarium graminearum | Nucleotide | 96.1 | 92.1 | 94.1 |
| geneML | Puccinia striiformis | Exon | 67.7 | 51.7 | 58.6 |
| geneML | Puccinia striiformis | Gene | 42.8 | 31.6 | 36.4 |
| geneML | Puccinia striiformis | Nucleotide | 89.7 | 62.8 | 73.9 |
| geneML | Rhizophagus irregularis | Exon | 68.6 | 51.2 | 58.6 |
| geneML | Rhizophagus irregularis | Gene | 50.4 | 35.8 | 41.9 |
| geneML | Rhizophagus irregularis | Nucleotide | 93.6 | 65 | 76.7 |
| geneML | Saccharomyces cerevisiae | Exon | 85.5 | 70.5 | 77.3 |
| geneML | Saccharomyces cerevisiae | Gene | 88 | 92.2 | 90.1 |
| geneML | Saccharomyces cerevisiae | Nucleotide | 98.2 | 98.5 | 98.3 |
| geneML | Somion occarium | Exon | 72.7 | 84.9 | 78.3 |

| | | | | | |
|---|---|---|---|---|---|
| geneML | Somion occarium | Gene | 70.1 | 67.5 | 68.8 |
| geneML | Somion occarium | Nucleotide | 95 | 91.8 | 93.4 |
| Helixer | Aspergillus flavus | Exon | 74.6 | 71.9 | 73.2 |
| Helixer | Aspergillus flavus | Gene | 56.6 | 56.1 | 56.3 |
| Helixer | Aspergillus flavus | Nucleotide | 97.4 | 95.6 | 96.5 |
| Helixer | Botrytis cinerea | Exon | 80.1 | 74.8 | 77.4 |
| Helixer | Botrytis cinerea | Gene | 67.2 | 61 | 64.0 |
| Helixer | Botrytis cinerea | Nucleotide | 99.1 | 94.2 | 96.6 |
| Helixer | Cercospora beticola | Exon | 77.6 | 66.2 | 71.4 |
| Helixer | Cercospora beticola | Gene | 63.1 | 55.9 | 59.3 |
| Helixer | Cercospora beticola | Nucleotide | 99.4 | 92.1 | 95.6 |
| Helixer | Cryptococcus tetragattii | Exon | 75 | 74.9 | 74.9 |
| Helixer | Cryptococcus tetragattii | Gene | 35.9 | 34.5 | 35.2 |
| Helixer | Cryptococcus tetragattii | Nucleotide | 98.1 | 95.6 | 96.8 |
| Helixer | Fusarium graminearum | Exon | 58.4 | 58.7 | 58.5 |
| Helixer | Fusarium graminearum | Gene | 42.7 | 43.2 | 42.9 |
| Helixer | Fusarium graminearum | Nucleotide | 97.1 | 91.3 | 94.1 |
| Helixer | Puccinia striiformis | Exon | 64.7 | 43.1 | 51.7 |

| | | | | | |
|---|---|---|---|---|---|
| Helixer | Puccinia striiformis | Gene | 28.4 | 19.4 | 23.1 |
| Helixer | Puccinia striiformis | Nucleotide | 93.2 | 61.5 | 74.1 |
| Helixer | Rhizophagus irregularis | Exon | 64.1 | 49.2 | 55.7 |
| Helixer | Rhizophagus irregularis | Gene | 39.6 | 29.3 | 33.7 |
| Helixer | Rhizophagus irregularis | Nucleotide | 95.3 | 66.6 | 78.4 |
| Helixer | Saccharomyces cerevisiae | Exon | 83.5 | 82.4 | 82.9 |
| Helixer | Saccharomyces cerevisiae | Gene | 85.1 | 88.6 | 86.8 |
| Helixer | Saccharomyces cerevisiae | Nucleotide | 98.7 | 98.7 | 98.7 |
| Helixer | Somion occarium | Exon | 66.3 | 77.5 | 71.5 |
| Helixer | Somion occarium | Gene | 45.6 | 42.6 | 44.0 |
| Helixer | Somion occarium | Nucleotide | 95.6 | 91.5 | 93.5 |

Table S7: Number of annotated genes with complete, partial or no PFAM domain(s) per tool, for the *Puccinia striiformis* and *Rhizophagus irregularis* reference genomes.

| genome | PFAM domains | Reference | ANNEVO | AUGUSTUS | BRAKER3 | Helixer | geneML |
|---|---|---|---|---|---|---|---|
| Puccinia striiformis | None | 9251 | 9622 | 3727 | 8578 | 14428 | 13069 |
| Puccinia striiformis | Partial | 5203 | 5152 | 1845 | 6289 | 6054 | 6280 |
| Puccinia striiformis | Complete | 2664 | 2830 | 1191 | 4219 | 4490 | 4579 |
| Rhizophagus irregularis | None | 18281 | 17830 | 6293 | 17916 | 22992 | 26139 |
| Rhizophagus irregularis | Partial | 7254 | 7265 | 2910 | 10625 | 10598 | 10553 |
| Rhizophagus irregularis | Complete | 4612 | 4046 | 1962 | 6719 | 7039 | 7241 |

Table S8: Runtime of different gene prediction tools per benchmarking genome, measured on a system with 8 available CPU cores

| Tool | Species | Runtime (hours) | Runtime (minutes) |
|---|---|---|---|
| ANNEVO | Aspergillus flavus | 0.21 | 12.8 |
| ANNEVO | Botrytis cinerea | 0.22 | 13.3 |
| ANNEVO | Cercospora beticola | 0.20 | 11.9 |
| ANNEVO | Cryptococcus tetragattii | 0.10 | 6.3 |
| ANNEVO | Fusarium graminearum | 0.21 | 12.7 |
| ANNEVO | Puccinia striiformis | 0.49 | 29.6 |
| ANNEVO | Rhizophagus irregularis | 0.78 | 46.9 |
| ANNEVO | Saccharomyces cerevisiae | 0.07 | 4.2 |
| ANNEVO | Somion occarium | 0.17 | 10.1 |
| AUGUSTUS | Aspergillus flavus | 0.65 | 39.3 |
| AUGUSTUS | Botrytis cinerea | 0.82 | 49.1 |
| AUGUSTUS | Cercospora beticola | 0.56 | 33.8 |
| AUGUSTUS | Cryptococcus tetragattii | 0.24 | 14.5 |
| AUGUSTUS | Fusarium graminearum | 0.62 | 37.1 |
| AUGUSTUS | Puccinia striiformis | 2.25 | 135.0 |
| AUGUSTUS | Rhizophagus irregularis | 3.48 | 209.0 |

| | | | |
|---|---|---|---|
| AUGUSTUS | Saccharomyces cerevisiae | 0.17 | 10.1 |
| AUGUSTUS | Somion occarium | 0.67 | 40.2 |
| BRAKER3 | Aspergillus flavus | 2.70 | 161.8 |
| BRAKER3 | Botrytis cinerea | 3.17 | 190.1 |
| BRAKER3 | Cercospora beticola | 2.45 | 146.8 |
| BRAKER3 | Cryptococcus tetragattii | 1.87 | 112.2 |
| BRAKER3 | Fusarium graminearum | 2.89 | 173.6 |
| BRAKER3 | Puccinia striiformis | 4.42 | 265.3 |
| BRAKER3 | Rhizophagus irregularis | 8.97 | 538.0 |
| BRAKER3 | Saccharomyces cerevisiae | 1.16 | 69.4 |
| BRAKER3 | Somion occarium | 2.47 | 148.2 |
| geneML | Aspergillus flavus | 0.06 | 3.7 |
| geneML | Botrytis cinerea | 0.05 | 3.3 |
| geneML | Cercospora beticola | 0.06 | 3.3 |
| geneML | Cryptococcus tetragattii | 0.07 | 4.4 |
| geneML | Fusarium graminearum | 0.07 | 4.2 |
| geneML | Puccinia striiformis | 0.19 | 11.6 |
| geneML | Rhizophagus irregularis | 0.20 | 12.3 |

| | | | |
|---|---|---|---|
| geneML | Saccharomyces cerevisiae | 0.02 | 1.4 |
| geneML | Somion occarium | 0.20 | 12.1 |
| Helixer | Aspergillus flavus | 0.14 | 8.3 |
| Helixer | Botrytis cinerea | 0.15 | 9.2 |
| Helixer | Cercospora beticola | 0.14 | 8.2 |
| Helixer | Cryptococcus tetragattii | 0.09 | 5.4 |
| Helixer | Fusarium graminearum | 0.14 | 8.2 |
| Helixer | Puccinia striiformis | 0.36 | 21.9 |
| Helixer | Rhizophagus irregularis | 0.54 | 32.3 |
| Helixer | Saccharomyces cerevisiae | 0.05 | 3.0 |
| Helixer | Somion occarium | 0.12 | 7.4 |

Table S9: Log-log regression values of runtime by genome size per benchmarked tool.

| Tool | a | b | r | R2 | p value |
|---|---|---|---|---|---|
| geneML | 0.01 | 0.72 | 0.74 | 0.54 | 2.33E-02 |
| BRAKER3 | 0.19 | 0.74 | 0.98 | 0.96 | 3.45E-06 |
| Helixer | 0.01 | 0.93 | 0.99 | 0.98 | 1.76E-07 |
| ANNEVO | 0.01 | 0.97 | 1.00 | 1.00 | 1.26E-10 |
| AUGUSTUS | 0.01 | 1.25 | 0.99 | 0.99 | 1.13E-07 |

Table S10: Statistics on the number of annotated genes per genome in the original annotation versus geneML annotation of training genomes (n=761).

| dataset | median | mean | min | max | q1 | q3 |
|---|---|---|---|---|---|---|
| geneML | 3547 | 4068 | 158 | 25457 | 2683 | 5032 |
| reference | 3241 | 3782 | 7 | 61481 | 2462 | 4837 |

Table S11: Total number of annotated genes with complete, partial or no PFAM domain(s) by phylum, for geneML annotations of training genomes (n=761), compared to original NCBI annotations.

| Phylum | Annotation dataset | Genes without PFAM domains | Genes with partial PFAM domains | Genes with complete PFAM domains |
|---|---|---|---|---|
| Ascomycota | original | 1563473 | 2340614 | 1214062 |
| Ascomycota | geneml | 1567728 | 2454779 | 1304687 |
| Basidiomycota | original | 1138353 | 917643 | 484512 |
| Basidiomycota | geneml | 1205539 | 1019157 | 629696 |
| Other | original | 357133 | 376634 | 241119 |
| Other | geneml | 410655 | 392905 | 302287 |
| All | original | 3058959 | 3634891 | 1939693 |
| All | geneml | 3183922 | 3866841 | 2236670 |

Table S12: Number of genes assigned to functional bins based on original NCBI annotations and geneML annotations of training genomes (n=761).

| Functional bin | original | geneML |
|---|---|---|
| Cell structure & cell cycle | 28495 | 30181 |
| General metabolism | 1159888 | 1211683 |
| Genetic information processing | 353612 | 376435 |
| Mobile genetic elements | 29784 | 130213 |
| Protein homeostasis & turnover | 76419 | 81012 |
| Regulation & signaling | 330580 | 376348 |
| Secondary metabolism | 100467 | 106877 |
| Transport & trafficking | 394255 | 414128 |

Table S13: PFAM domains significantly overrepresented in geneML annotations compared to original NCBI annotations. Showing the top 20 domains with the largest absolute increase in gene counts.

| Bin ID | PFAM ID | PFAM description | n paired genomes | median original | median geneML | total diff (geneml-original) | p value | q value |
|---|---|---|---|---|---|---|---|---|
| 1 | PF00078.33 | Reverse transcriptase (RNA-dependent DNA polymerase) | 740 | 1 | 10 | 47137 | 1.35E-94 | 1.12E-90 |
| 1 | PF17917.8 | RNase H-like domain found in reverse transcriptase | 601 | 0 | 11 | 37313 | 1.93E-83 | 4.81E-80 |
| 1 | PF17919.8 | RNase H-like domain found in reverse transcriptase | 597 | 0 | 12 | 37263 | 9.56E-83 | 2.16E-79 |
| 1 | PF17921.8 | Integrase zinc binding domain | 733 | 1 | 6 | 33667 | 1.06E-87 | 3.30E-84 |
| 1 | PF24626.2 | Tf2-1-like, SH3 domain | 550 | 0 | 9 | 27332 | 2.11E-80 | 4.38E-77 |
| 3 | PF00172.24 | Fungal Zn(2)-Cys(6) binuclear cluster domain | 751 | 69 | 96 | 24545 | 3.76E-106 | 9.35E-102 |
| 1 | PF00665.33 | Integrase core domain | 492 | 0 | 12.5 | 23160 | 2.34E-71 | 2.78E-68 |

| | | | | | | | | |
|---|---|---|---|---|---|---|---|---|
| 1 | PF00385.31 | Chromo (CHRromatin Organisation MOdifier) domain | 490 | 0 | 11 | 21086 | 4.48E-73 | 6.57E-70 |
| 1 | PF30863.1 | PEG10/RTL1 protease domain | 418 | 0 | 9 | 16938 | 4.20E-63 | 2.99E-60 |
| 1 | PF03184.26 | DDE superfamily endonuclease | 578 | 0 | 6 | 13827 | 4.90E-74 | 8.13E-71 |
| 3 | PF04082.24 | Fungal specific transcription factor domain | 748 | 35 | 46 | 11895 | 5.02E-102 | 6.25E-98 |
| 4 | PF00098.29 | Zinc knuckle | 758 | 6 | 9 | 9088 | 3.99E-87 | 1.10E-83 |
| 1 | PF08284.17 | Retroviral aspartyl protease | 320 | 0 | 8 | 8964 | 3.17E-47 | 1.43E-44 |
| 6 | PF07690.22 | Major Facilitator Superfamily | 761 | 176 | 189 | 8616 | 1.80E-53 | 9.98E-51 |
| 3 | PF00069.32 | Protein kinase domain | 761 | 126 | 130 | 7743 | 8.15E-63 | 5.49E-60 |
| 3 | PF07714.24 | Protein tyrosine and serine/threonine kinase | 761 | 114 | 119 | 7064 | 1.09E-65 | 1.01E-62 |
| 1 | PF03221.23 | Tc5 transposase DNA-binding domain | 453 | 0 | 3 | 6046 | 2.59E-63 | 1.89E-60 |
| 1 | PF01498.25 | Transposase | 371 | 0 | 4 | 5545 | 1.42E-57 | 8.42E-55 |

| | | | | | | | | |
|---|---|---|---|---|---|---|---|---|
| 1 | PF22936.2 | Pol polyprotein, beta-barrel domain | 287 | 0 | 7 | 5354 | 5.02E-44 | 1.87E-41 |
| 1 | PF07727.20 | Reverse transcriptase (RNA-dependent DNA polymerase) | 300 | 0 | 5 | 5283 | 5.72E-44 | 2.10E-41 |

### Supplementary Figures

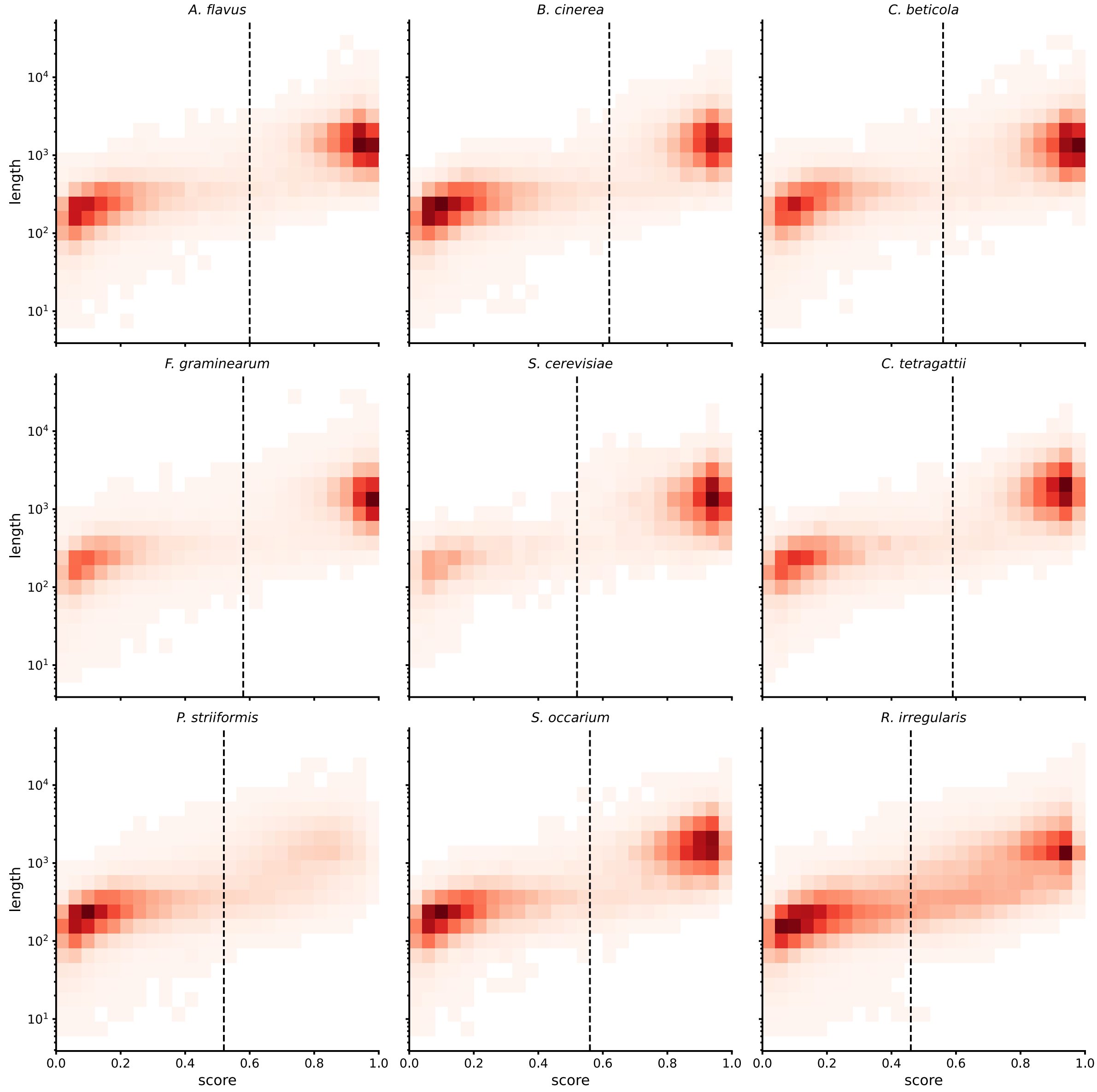


Figure S1: Distribution of geneML predicted genes by gene length and gene score, for each benchmarking genome. The chosen score threshold for each genome is shown as a dotted line.

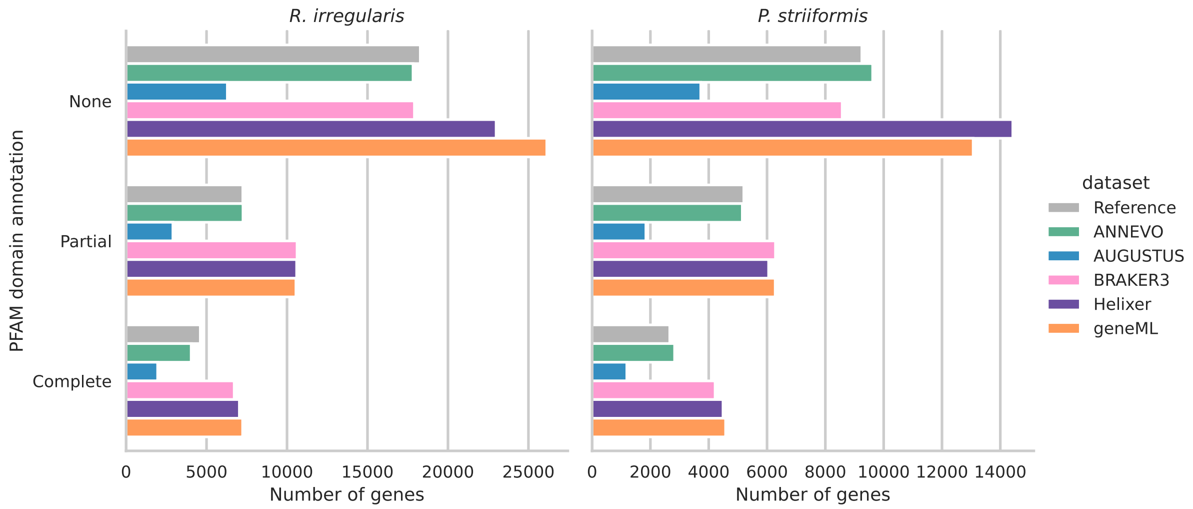


Figure S2: Number of annotated genes with complete, partial or no PFAM domain(s) per tool, for the *Puccinia striiformis* and *Rhizophagus irregularis* reference genomes.

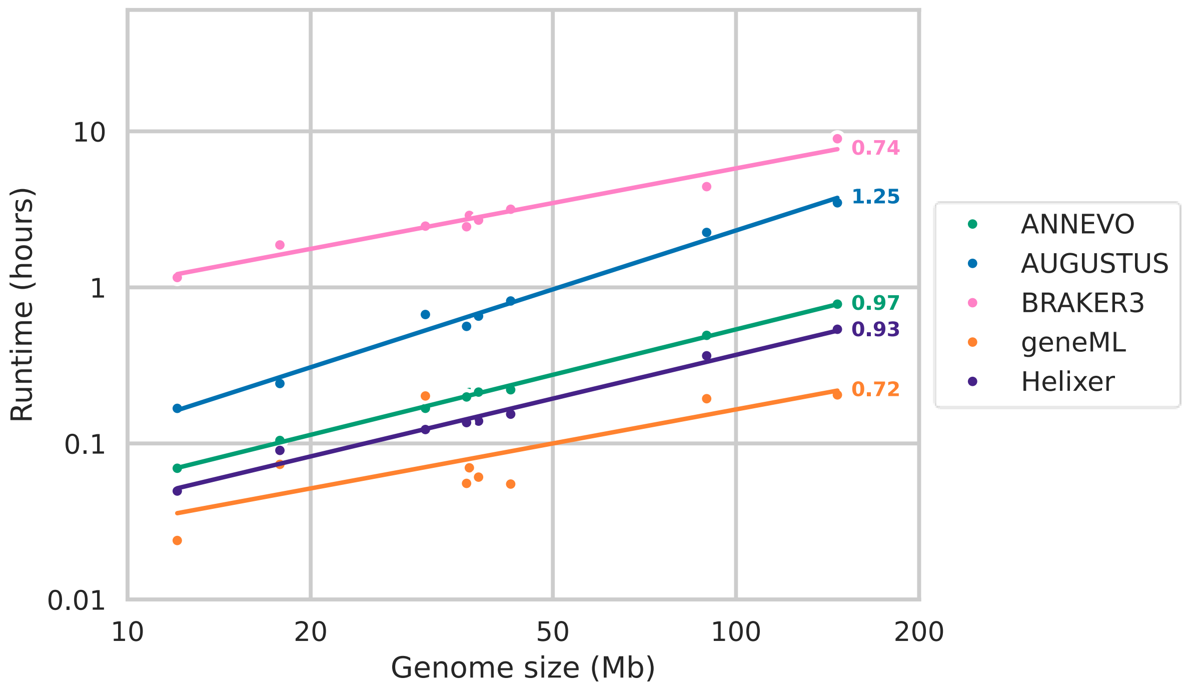


Figure S3: Runtime of different gene prediction tools by benchmarking genome size. For each tool, a log-log regression line and its slope are shown.

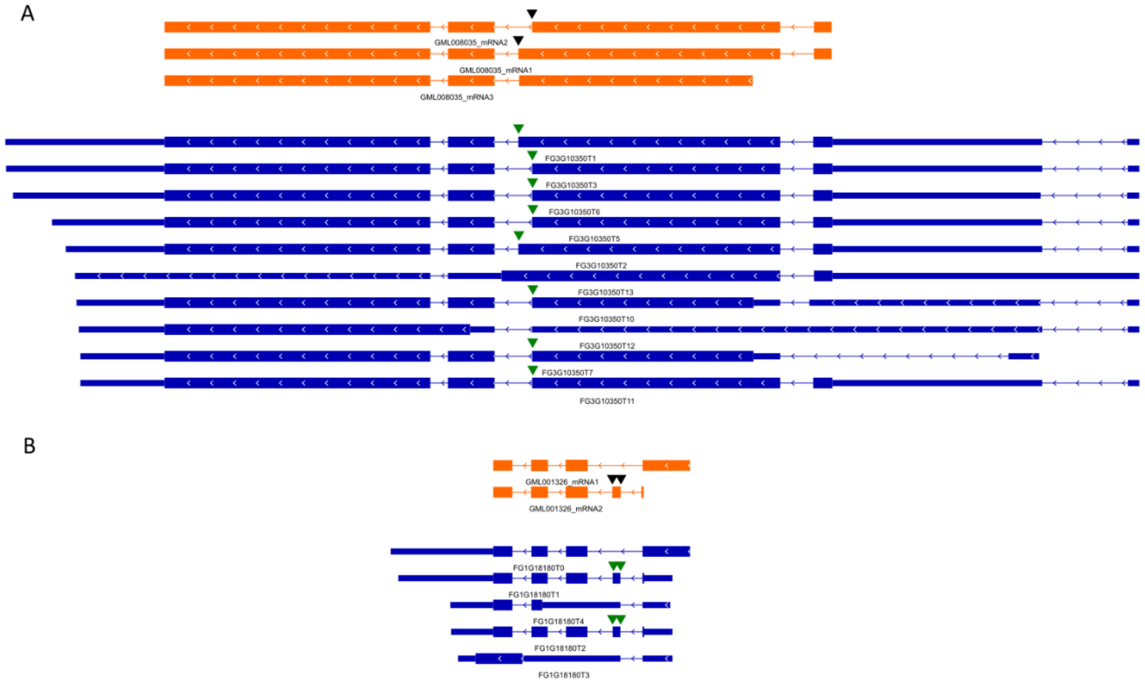


Figure S4: Additional examples of alternative transcript predictions supported by IsoSeq sequencing data: A) alternative 5’ splice site (10/15 reference transcripts shown), B) exon skipping. Transcripts predicted by geneML are shown in orange (predictions do not include UTRs), reference transcripts are shown in dark blue. Gene structures follow IGV conventions: thin lines = introns, medium bars = UTRs, thick bars = coding exons. Black inverted triangles indicate locations of predicted transcript variation, green inverted triangles indicate matching reference variants. *Fusarium graminearum* PH-1 transcript annotations were downloaded from http://fgbase.wheatscab.com/Downloadexon skipping and alternative 5’ splice site.

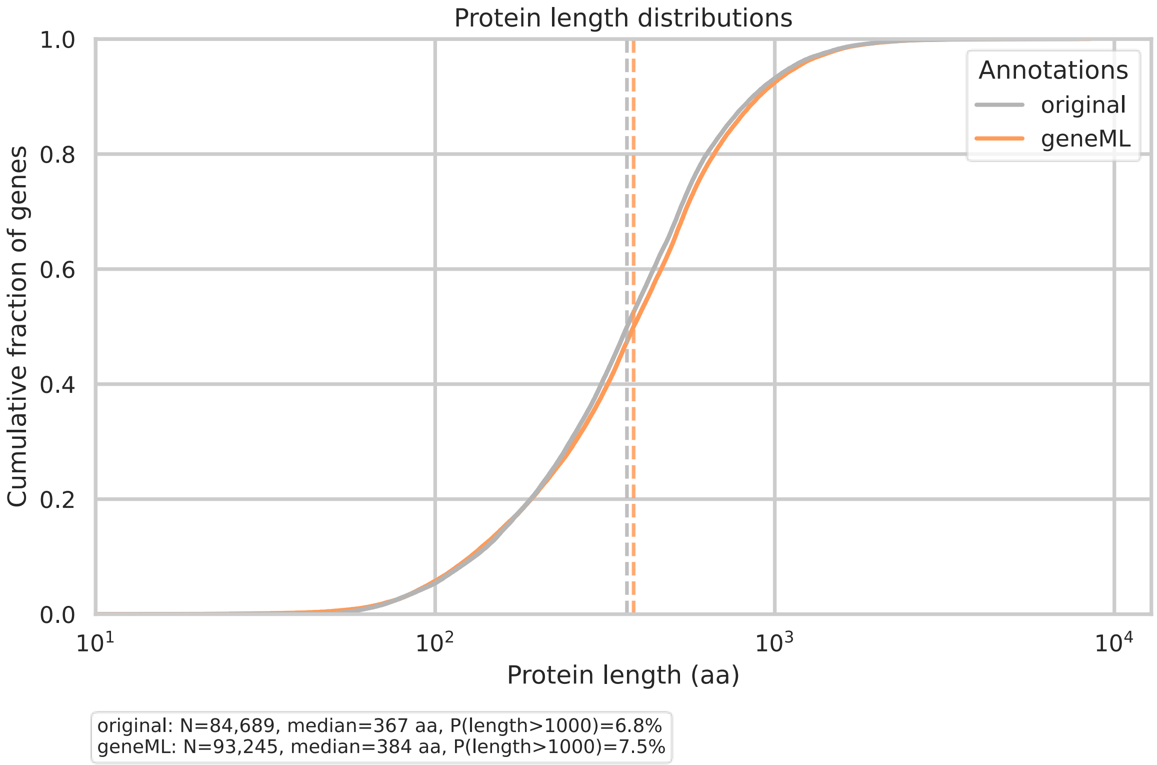


Figure S5: Distribution of protein lengths for geneML annotations versus original NCBI annotations across training genomes (n=761). Data were subsampled to every 100th protein, and only the longest isoform per locus was included.

#### Statement on the use of Large Language Models

Github Copilot was used to provide code suggestions and Anthropic Claude was used to suggest improvements to the grammar and clarity of written text. The authors take full responsibility for the final code and manuscript.
